# Localization of Macromolecules in Crowded Cellular Cryo-electron Tomograms from Extremely Sparse Labels

**DOI:** 10.1101/2024.11.04.620735

**Authors:** Mostofa Rafid Uddin, Ajmain Yasar Ahmed, H.M. Shadman Tabib, Md Toki Tahmid, Md Zarif Ul Alam, Zachary Freyberg, Min Xu

## Abstract

**Motivation:** Localizing macromolecules in crowded cellular cryo-electron tomograms (cryo-ET) is crucial for determining their *in situ* structures. Traditional template matching-based approaches for this task suffer from template-specific biases and have low throughput. Given these problems, learning-based solutions are necessary. However, the paucity of annotated data for training poses substantial challenges for such learning-based methods. Moreover, preparing extensively annotated cellular cryo-ET tomograms for training macromolecule localization methods is extremely time-consuming and burdensome due to the large volume and low signal-to-noise ratio of the tomograms.

**Results:** In this work, we developed TomoPicker, an annotation-efficient macromolecule localization method for cryo-ET tomograms. To achieve such annotation-efficiency, TomoPicker regards macromolecule localization as a voxel classification problem and solves it with two different positive-unlabeled learning approaches. We evaluated TomoPicker on two experimental cryo-electron tomography (cryo-ET) datasets of crowded eukaryotic cells and one experimental dataset of relatively less crowded prokaryotic cell. We observed that, with only 10 annotated macromolecule locations, TomoPicker with positive unlabeled learning achieved a performance comparable to that of state-of-the-art supervised methods trained with several hundred annotations. In other words, TomoPicker achieved plausible segmentation with up to 98% less data compared to supervised learning-based methods. Furthermore, it demonstrated substantial improvements over existing learning-based macromolecule localization methods under sparse annotation scenarios.

**Code:** The code to train and use TomoPicker is available on https://github.com/DuranRafid/TomoPicker.

## Introduction

Cellular cryo-electron tomography (Cryo-ET) is a prominent imaging technology that has enabled *in situ* 3D visualization of macromolecules up to subnanometer resolution within cells in near-native context [4, 17]. Furthermore, cryo-ET can resolve the structures of macromolecules within cells with different compositions and conformations. In addition, mapping macromolecules back to their original positions within cryo-electron tomograms can reveal the spatial organization of macromolecules within cells, providing potentially novel biological insights [14]. As a result, cryo-ET is emerging as a powerful imaging approach for *in situ* structural biology.

Determining the *in situ* structures of macromolecules from 3D cellular cryo-ET tomograms is a complex process involving multiple steps [2, 17]. The first and most crucial step is to localize the macromolecules in the tomograms [15, 8]. This is a challenging task for several reasons. Firstly, cellular cryo-ET tomograms are large 3D volumes with a size of approximately 1000 × 1000 × 500 pixels, even after 4 times of binning [19, 11]. Secondly, due to the complex cytoplasmic environment and the low electron dose to prevent radiation damage to the cellular specimen, these tomograms have a very low signal-to-noise ratio and contrast[11, 2]. Finally, the concentration of macromolecules per image is very high (∼ 500 − 1000 per cryo-tomogram) [8], making it even more challenging to locate them accurately.

Given the above-mentioned challenges, manually localizing macromolecules in the tomograms is extremely tedious and burdensome. To this end, automated approaches for macromolecule localization have been developed [15, 18, 12]. A common approach is template matching (TM), which uses templates from existing data sources, such as PDB, EMDB, etc., as references to localize similar macromolecules in the tomograms [1]. However, TM can only be applied when a reference template is available for the macromolecules to be localized and often contains reference-dependent biases [19, 11]. In addition, TM is extremely time-consuming [19] and shows suboptimal performance [10]. To solve this issue, neural network-based deep learning methods have been introduced [3, 10, 12, 18]. These methods provide high-throughput, fast localization of macromolecules without reference-dependent biases. However, most of these methods [12, 18] are based on supervised learning, which again requires manual annotation of a large number of macromolecule locations in the tomograms for training. Given the difficulty of manual annotation in cryo-ET tomograms, annotation-efficient methods that can perform reliable localization of macromolecules without requiring large annotated training data are necessary.

In recent years, a few learning-based macromolecule localization methods have addressed this annotation burden [5, 10]. Huang et al. [5] developed an algorithm to localize macromolecules from sparse labels by regarding the localization as a regression problem. The algorithm regards every 3D tomogram as a single sample and thus predicts macromolecule coordinates directly at the tomogram level. This approach has two issues. First, since this method is a learning-based method and regards each tomogram as a sample, a large number of similar tomograms are required in the training dataset. Second, fitting tomograms as inputs to convolutional networks requires significant downsampling of the tomograms. This downsampling exacerbates the existing macromolecular crowding within the tomograms, further reducing the signal-to-noise ratio. Such extra crowding makes this approach less suitable to highly crowded eukaryote cell tomograms. Another supervised algorithm has been developed very recently, called DeepETPicker [10]. Unlike the algorithm developed by Huang et al. [5], DeepETPicker can be trained on a single tomogram where several macromolecule coordinates in the tomogram are annotated. It works by placing sphere in annotated regions, extracting subvolumes with spheres, and training a supervised segmentation model with dice loss on the extracted subvolumes. Despite achieving success for several sparse single-particle and prokaryotic tomograms, its efficacy in crowded eukaryotic tomograms has not been explored. Moreover, DeepETPicker [10] did not adapt any specific mechanism in its training process to deal with the annotation-efficiency issue. Hence, its performance substantially declines under extremely sparse annotations (Fig. 2, 3, 5). CryoSAM [20] is also a recently developed method that uses a small amount of annotated macromolecules as prompts to localize other macromolecules in a tomogram with vision foundation models. Despite demonstrating some success in localizing macromolecules from specifically denoised sparse prokaryote tomograms, its efficacy in crowded eukaryotic tomograms or under extremely sparse label conditions is yet to be examined. Consequently, an annotation-efficient approach for macromolecule localization in crowded cellular cryo-ET tomograms under extremely sparse labels is needed.

**Fig. 1.**
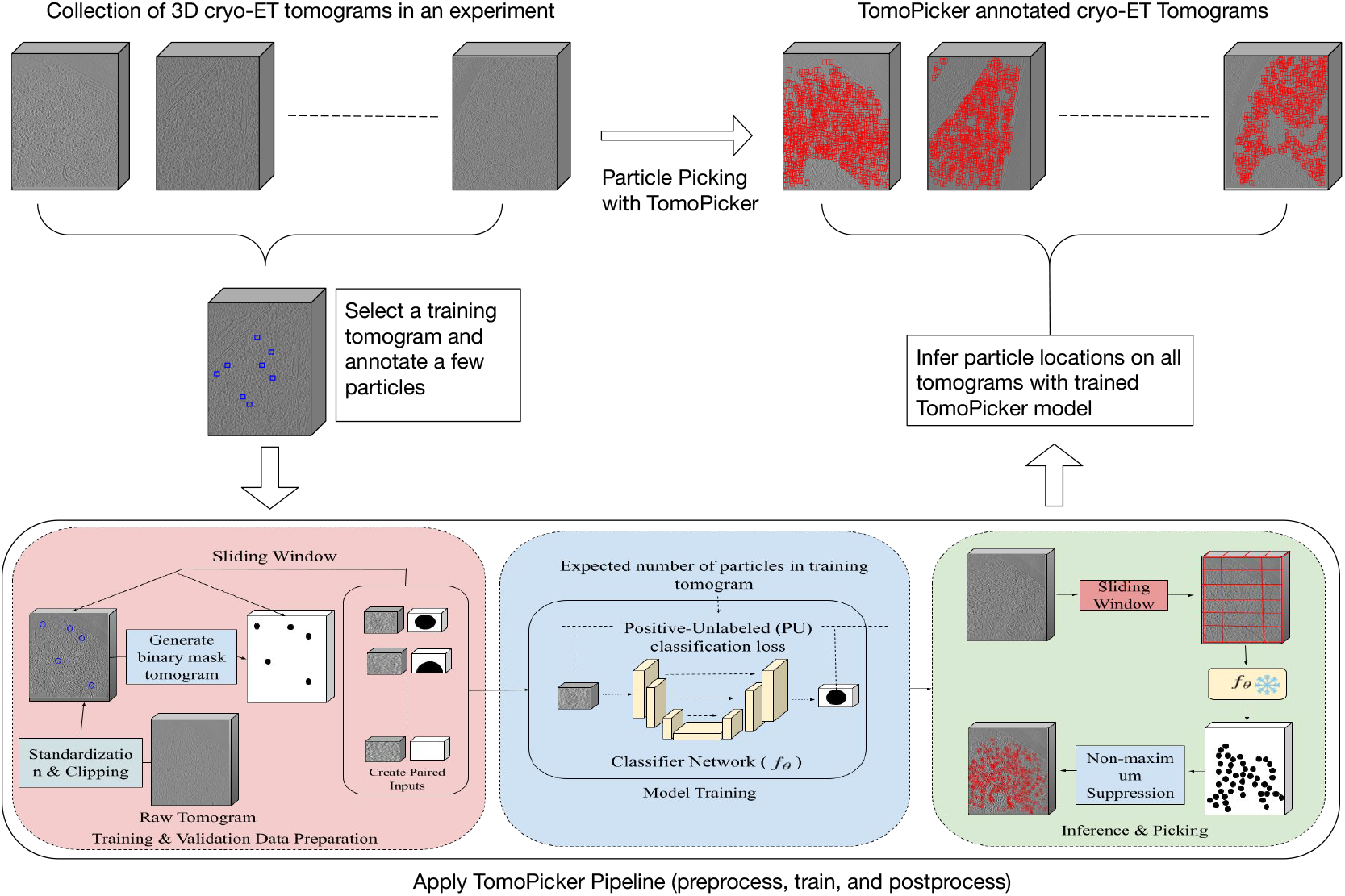
Overview of the macromolecule localization process with TomoPicker.

**Fig. 2.**
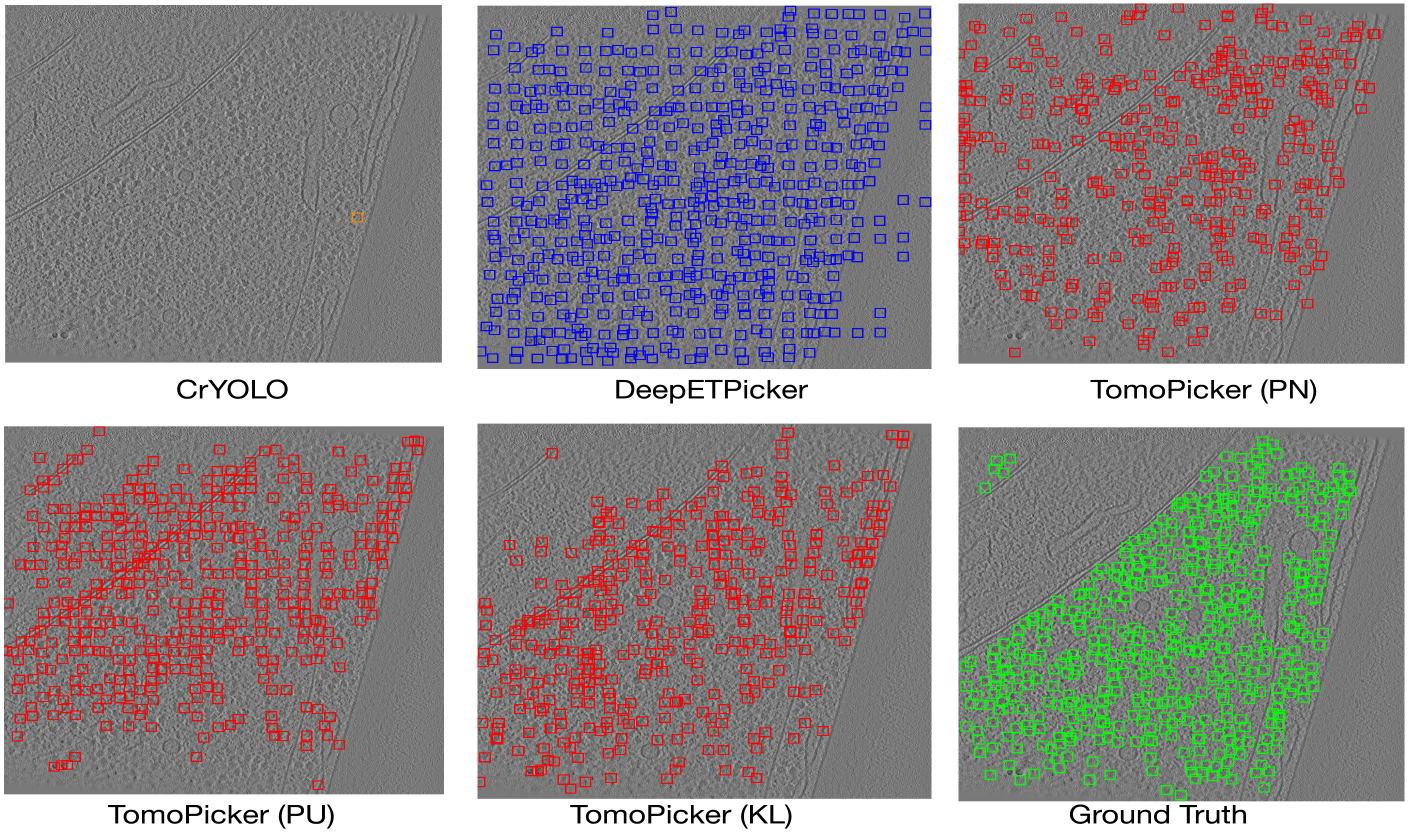
Predictions vs Ground Truth for central (*z* = 500) xy slice of TS 0003 VPP tomogram. Green Box = Ground Truth, Red Box = TomoPicker Predictions, Blue Box = DeepETPicker Predictions, Orange Box = CrYOLO Predictions.

**Fig. 3.**
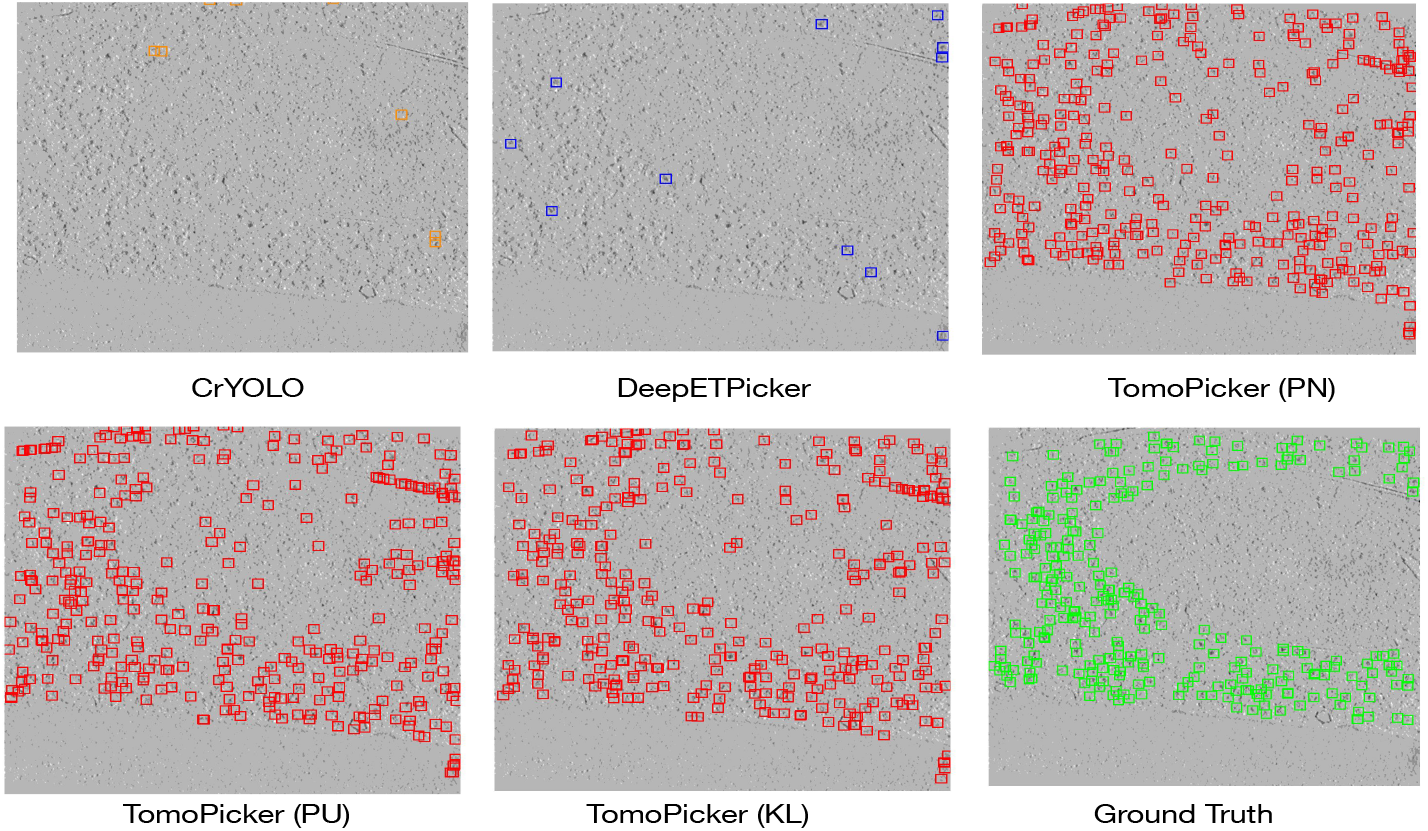
Predictions vs Ground Truth for central (*z* = 250) xy slice of denoised TS 045 defocus-only tomogram. Green Box = Ground Truth, Red Box = TomoPicker Predictions, Blue Box = DeepETPicker Predictions, Orange Box = CrYOLO Predictions.

**Fig. 4.**
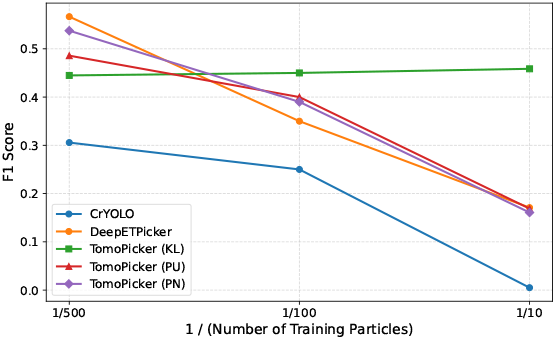
Average F1 score of TomoPicker with different losses (PN, PU, KL), DeepETPicker [10], and CrYOLO [18] on VPP *S.Pombe* dataset with varying number of macromolecular structures used for training

**Fig. 5.**
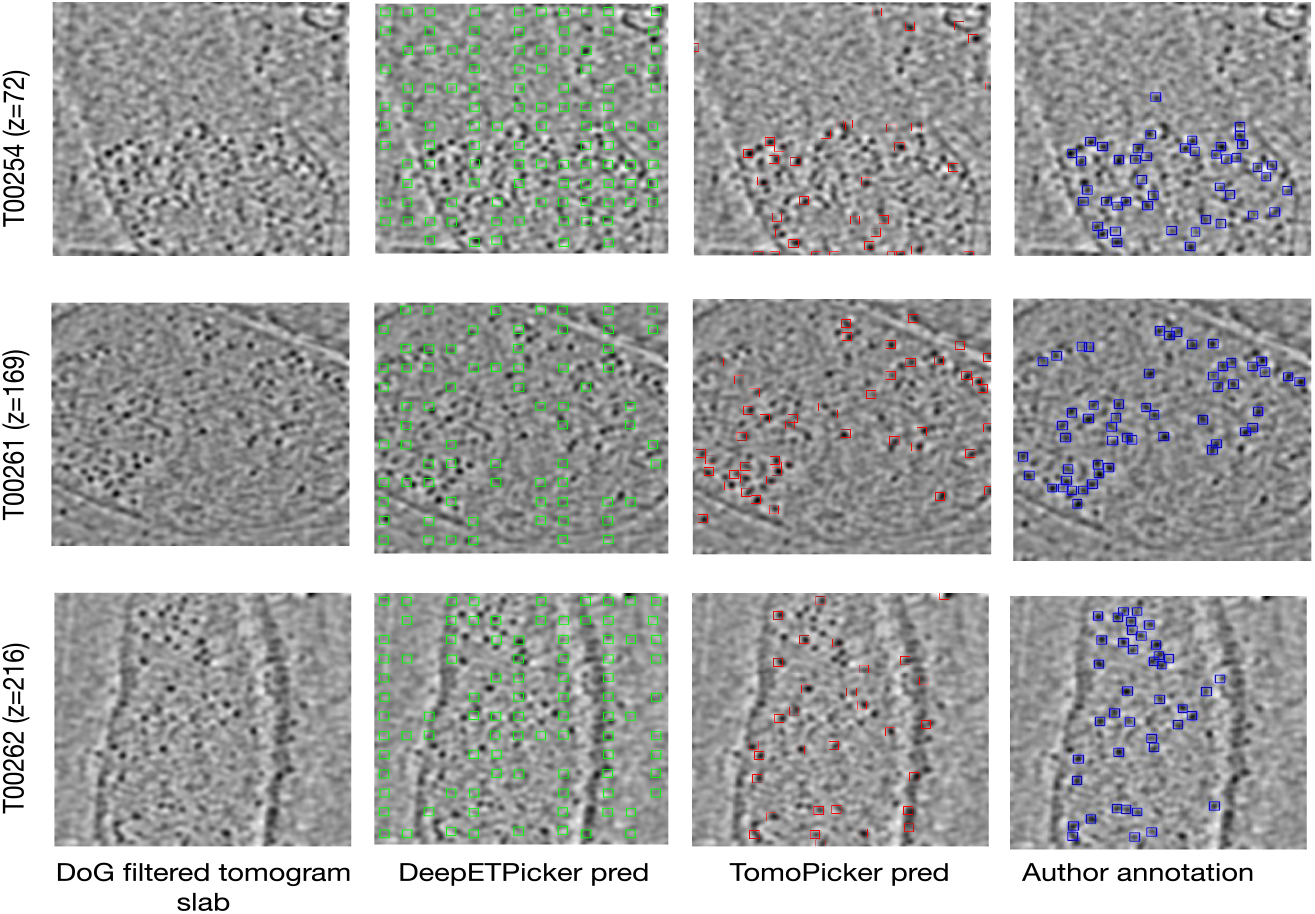
Macromolecule localization results by TomoPicker (KL) and DeepETPicker (both trained using 10 annotated locations) on *M. Pneumoniae* cellular cryo-ET tomograms from EMPIAR-10499. The figure shows three slabs from three different tomograms in the dataset (after DoG filtering for better visualization), the inference results by DeepETPicker and TomoPicker (KL), and the author annotations in the corresponding slabs.

In this work, we developed a novel annotation-efficient macromolecule localization approach called TomoPicker. Our approach requires only 10 macromolecule locations from a single training tomogram to be annotated beforehand. In TomoPicker, we regard macromolecule localization as a voxel classification problem. For 3D cryo-ET tomograms, each voxel is classified as a binary value based on whether it contains macromolecules or not. Given that only a few voxels are labeled as positive based on manual annotation, a specific approach is necessary to deal with the large number of unlabeled voxels. If all unlabeled voxels are regarded as negative, it would lead to erroneous prediction and macromolecule localization. To solve this problem, we introduce two positive-unlabeled (PU) learning approaches-one based on non-negative risk estimation and another based on regularization based on a prior distribution, with the latter being our primary method.

We primarily evaluated TomoPicker against two well-annotated benchmark datasets of eukaryotic *S. Pombe* cellular cryo-ET tomograms, where one of the datasets is imaged with Volta-Phase-Plate (VPP) and the other is imaged without it. We also evaluated it against a well-known prokaryotic *M. Pneumoniae* cellular cryo-ET dataset. Along with TomoPicker, we tested recent and popular learning-based cryoET macromolecule localization methods (including the state-of-the-art DeepETPicker [10]) in these data sets. Our extensive experiments demonstrate superior performance of the TomoPicker approach compared to other learning-based methods. Using only 10 macromolecule locations as manual annotation, which is 0.04% of the total number of macromolecules, TomoPicker provides predictions that most closely resemble the expert annotations (Fig. 2 and Fig. 3). In addition, TomoPicker achieves the performance of state-of-the-art macromolecule localization methods with 98% less annotation. Under extremely sparse annotation scenarios, it substantially outperforms the other methods. Overall, our experiments suggest that TomoPicker is a valuable tool for macromolecule localization in 3D cellular cryo-ET tomograms, particularly in crowded cellular tomograms, in a highly annotation-efficient manner.

## Methods

Given a set of *n* ∈ ℕ tomograms 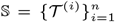, where each tomogram 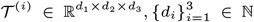 is a 3D grayscale image, TomoPicker provides a set of particle coordinates 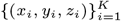 for each tomogram 𝒯^(*i*)^

TomoPicker pipeline consists of three main components (Fig. 1). We briefly discuss them as follows:

### Preprocessing and Data Generation for Training

The tomograms 𝒯 ^(*i*)^ in the dataset 𝕊 are preprocessed to enhance contrast. We load the tomograms as voxelized arrays and standardize the voxels for each tomogram to 0 mean and 1 standard deviation. Then, we clip those values that lie beyond three standard deviations from the mean (which is 0 after standardization). After clipping, we again standardized the voxels in each tomogram to 0 mean and 1 standard deviation.

A training dataset 𝕋 and validation dataset 𝕍 (optional) is selected from the dataset 𝕊, where |𝕋| ≪ |𝕊|. In our experiments, we used only one tomogram for training and one for validation, keeping |𝕋| = |𝕍| = 1. For each training and validation tomograms 𝒯 ^(*i*)^ ∈ {𝕋, 𝕍}, we create empty voxel arrays {ℒ^(*i*)^} having the same shape as the corresponding tomogram. We create these empty voxel arrays to generate labels for each voxel for our voxel classification-based macromolecule localization network. We require a few (minuscule percent of the total particles *K*) particles coordinates 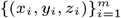, *m* ≪ *K* to be provided for the training and validation tomograms. For each of the provided particle coordinates per training or validation tomogram, we put values of 1 in all the voxels around the coordinate that lie within a distance equal to the radius *r* ∈ ℕ of that particle in the empty voxel arrays. Thus, we have a roughly spherical mask of 1s for each labeled particle in the corresponding {ℒ^(*i*)^} arrays belonging to the tomogram (Fig. 1).

Given the large size of the tomograms, it is difficult to pass them directly as input to our macromolecule-localization model. Consequently, we generate small subvolumes and submasks from the tomograms and their corresponding label arrays. To this end, we use a sliding window approach to extract subvolumes *t*^(*i*)^ ∈ ℝ^*s×s×s*^ and their corresponding submasks *l*^(*i*)^ ∈ {0, 1}^*s×s×s*^ with a given subtomogram size *s* from the tomogram 𝒯 ^(*i*)^ and label array ℒ^(*i*)^, respectively. Thus, we collect subvolumes and submask pairs (*t*^(*i*)^, *l*^(*i*)^) from the tomograms. We save all such pairs for training tomograms and use them to train our macromolecule localization model. For validation tomograms, we only save those pairs with submasks *l*^(*i*)^ with at least one non-zero value. After saving the data necessary for training and validation, we move forward to the next step of model training.

### Training Classifier with Positive Unlabeled (PU) Leaning

We formulate macromolecule localization as a voxel classification problem. Our training dataset consists of *N* number of (*t*^(*i*)^, *l*^(*i*)^) pairs where *l*^(*i*)^ serves as the voxel-wise label for the subvolume *t*^(*i*)^. We assume that *P*_*i*_ is the set of voxels in *t*^(*i*)^ that are labeled as 1 and *U*_*i*_ is the set of voxels in *t*^(*i*)^ labeled as 0. In other words, *P*_*i*_ is the set of voxels in a subtomogram that contains an annotated macromolecule, and *U*_*i*_ is the set of remaining voxels that contain both unlabeled macromolecules and non-particle regions in the training dataset. Given 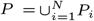 and 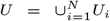, we learn a classifier (*f*_*θ*_) that distinguishes between macromolecule and non-macromolecule regions in the subtomograms. In other words, the classifier is our macromolecule localization model. We used three different strategies (two with PU learning and one without) to train the classifier resulting in three different methods, which we discuss below.

### Positive Negative (PN) Learning

When all voxels of *U* are from non-macromolecule regions, we can treat the voxels in *P* as positive voxels and all the voxels in *U* as negative voxels. In such a scenario, we train a classifier with a standard loss minimization objective function (named PN) as shown in Equation 1.

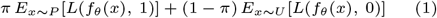

Here, *x* is a single voxel belonging to either *P* or *U*. *π* is the fraction of labeled and unlabeled macromolecule regions within *P* ∪ *U*. The value of *π* can be calculated as the fraction of non-zero values in the training data after the label generation process mentioned before. In other words,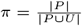.

We denote *L* as the cost function between the classifier output and labels. To implement it, we use binary cross-entropy loss defined as follows:

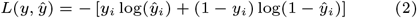

Here, *ŷ* is the prediction *f*_*θ*_(*x*) and *y* is the ground truth label from *l*^(*i*)^.

We refer to the TomoPicker method trained with this PN learning objective as TomoPicker (PN).

However, this PN objective function is effective when all the macromolecule regions in tomograms are labeled, thus belonging only to *P*, and only non-macromolecule regions are in *U*, which is usually not the case. In an ideal scenario, *P* contains voxels of only a few macromolecules in the tomogram, whereas most of the macromolecule voxels belong in *U*. As a result, PN learning provides suboptimal macromolecule localization performance in our experiments. To this end, we introduce positive-unlabeled (PU) learning.

### Non-negative Risk Estimator based Positive Unlabeled (PU) Leaning

We first leveraged a non-negative risk estimator-based PU Learning approach [9] to incorporate PU learning in our cryo-ET macromolecule localization framework. In this framework, we use *π*^*′*^ as the expected average of voxel values in the label of a voxel in the training tomogram. We calculate this based on the expected number of macromolecules per training tomogram, *Ep*, and the macromolecule radius *r* provided by the user. Based on *r*, a fixed number of voxels are labeled as 1 in the training data generation process. If this number is denoted as *Rn*, then the value of *π*^*′*^ is: 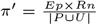. Here, *Ep* × *Rn* is the expected total number of 1s in |*P* ∩ *U* |.

We further denote 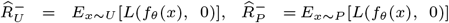, and 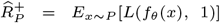. *L* is implemented as Equation 2. Using these three, we denote a new quantity 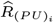 as 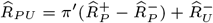

We used a specialized version of the algorithm in [9] to train our macromolecule localization network. If 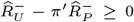, we update the parameters of *f*_*θ*_ using 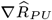, where ∇ is the gradient with respect to the parameters. Otherwise, we update the parameters of *f*_*θ*_ using 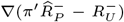. We refer to the TomoPicker method trained with this algorithm as TomoPicker (PU).

### KL based Positive Unlabeled Leaning Loss

We finally propose an alternate approach to non-negative risk estimator PU learning. Instead of minimizing the positive-negative misclassification loss, a classifier can simultaneously attempt to minimize the *P* class misclassification loss and match the expectation over *U*. In other words, we can learn a classifier (*f*_*θ*_) that minimizes *E*_*x*∼*P*_ [*L*(*f*_*θ*_(*x*), 1)] subject to the constraint *E*_*x*∼*U*_ [*f*_*θ*_(*x*)] = *π*^*»*^, where *π*^*”*^ is the fraction of voxels from unlabeled macromolecule regions within *U*.

We can impose such a constraint through a regularization term in the objective function with a weight *λ* (set as 10 in our experiments) as shown in Equation 3.

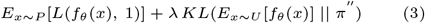

This objective imposes a constraint through the KL-divergence between the expectation of the classifier over *U* and the estimated fraction of voxels from unlabeled macromolecule regions in *U*, denoted by *π′′*. This divergence is minimized when log both terms are close to each other. KL divergence between two distributions *P* and *Q* can be defined as: 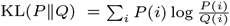.

To calculate *π′′*, we take the expected number of macromolecules per training tomogram (*Ep*) as input, similar to our previous non-negative risk estimator-based approach. Given the number of voxels *Rn* labeled as 1 in data generation for a macromolecule of radius *r*, the expected total 1 in *P* ∪ *U* is *Ep* × *Rn*. Among them, |*P* | is the observed number of 1. So, the expected number of unlabeled 1s that are unlabeled is *Ep* × *Rn* − |*P* |. So, *π′′* can be written as.

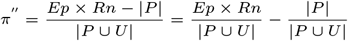

Using the definition of *π* and *π*^*′*^, *π′′* can be expressed as: *π′′* = *π*^*′*^ − *π*

We refer to the TomoPicker method trained with this KL-based objective as TomoPicker (KL).

### Implementation of the classifier

To implement the classifier network *f*_*θ*_, we used a MultiResUnet architecture [6]. Our MultiResUnet network consists of four down-sampling and four up-sampling layers. Each of the down-sampling layers consist of either ResUNet or UNet blocks, starting with an input channel of 1 and progressively increasing to 32, 64, 128, and 256 channels. The primary difference between a ResUNet block and UNet block is the use of a residual connection in the ResUNet block which adds the block’s input to its output. The UNet block, on the other hand, simply stacks two convolutional layers without any residual addition. The upsampling path mirrors this configuration, restoring spatial dimensions with transposed convolutions. Skip connections provide concatenation between the corresponding up-sampling and down-sampling layers. An optional dropout layer is used for regularization, and a final 1×1 3D convolution produces the single-channel output. All the experimental results presented here use the default configuration of using the ResUNet layer in the up-sampling and down-sampling networks with no additional dropout for regularization.

We train the classifier network using the losses mentioned above.—We use a mini-batch size *bs* of 8 and use Adam optimizer to optimize the parameters *θ* in *f*_*θ*_.

### Inference and Picking

After training the classifier with the above-mentioned learning strategies, we perform macromolecule localization on all the tomograms in the dataset, including the ones we used for training and validation. For each tomogram *V*, we use a sliding window strategy to obtain non-overlapping subvolumes of the same size as the training subtomograms. Then, we perform inference for each subvolume with our learned classifier *f*_*θ*_. The inference results in a score for each voxel in the subvolumes. We merge the score outputs for each subvolume in the tomogram to a volumetric array (*V*_score_) with the same size as the tomogram. We then apply the picking process on this merged predicted array *V*_score_. The process takes the required number of macromolecules *N* or subvolume score threshold *t* and the macromolecule radius *r* as input. It operates in 4 steps. In step 1, Find the point (*x*_max_, *y*_max_, *z*_max_) with maximum score value in *V*_score_. In step 2, we append (*x*_max_, *y*_max_, *z*_max_) as well as the score *V*_score_(*x*_max_, *y*_max_, *z*_max_) to the extracted macromolecule list. In step 3, we remove a roughly spherical region of macromolecule radius *r* around (*x*_max_, *y*_max_, *z*_max_) in *V*_score_ by setting their scores to −∞. This ensures that the same macromolecule will not be extracted more than once. Finally, we repeat steps 1 − 3 until *N* macromolecules are extracted or no prediction scores above the threshold *t* remain.

## Experiments & Results

### Benchmarking

For benchmarking, we primarily used well-annotated cryo-ET datasets of *S. Pombe* cells publicly available at EMPIAR-10988 [7], which represent some of the only available well-annotated tomograms in the cellular cryo-ET domain. It contains a volta-phase-plate (VPP) tomogram dataset and a defocus-only tomogram dataset of *S. Pombe* cell sections (voxel spacing 1.348 nm). We used both datasets in our experiments for benchmarking. We have also used the cellular cryo-ET tomograms of *M. Pneumoniae* bacteria from EMPIAR-10499 [16]. It contains binned tomograms of *M. Pneumoniae* cell sections with a voxel spacing of 1 nm.

The VPP dataset of *S. Pombe* cells contain 10 tomograms (labeled from TS 0001 to TS 0010 consecutively) with a total of 25, 311 80S ribosome macromolecules. The individual tomograms from TS 0001 to TS 0010 contains 2450, 2342, 2429, 2967, 3571, 1336, 617, 2744, 3482, and 3373 ribosome macromolecules respectively.

The defocus-only dataset of *S. Pombe* cells consist of 10 tomograms (labeled as TS 026, TS 027, TS 028, TS 029, TS 030, TS 034, TS 037, TS 041, TS 043, TS 045) with a total of 25, 901 80S ribosome macromolecules. The individual tomograms on the abovementioned sequence contains 838, 1673, 5305, 2897, 2783, 3783, 1646, 2813, 1815, and 2348 ribosome macromolecules respectively. Since no volta-phase plate was used while capturing the cryo-ET images for the tomograms, the signal-to-noise ratio is much lower for these tomograms compared to the VPP tomograms. To this end, we denoised the tomograms with the CCP-EM denoiser [13] before training TomoPicker and the baseline methods.

The dataset of *M. Pneumoniae* cells consist of 65 tomograms of size 306 × 630 × 630. The exact number of ribosomes in these tomograms is unknown. However, several hundreds annotated ribosomes per tomogram are available from CZI cryo-ET data portal with high confidence of annotation. To train our method and the baselines, we used 10 samples from these annotations.

### Baselines

We used CrYOLO [18] and DeepETPicker [10] learning-based cryo-ET macromolecule localization methods as baselines with the aim of localizing ribosomal macromolecules. Since CrYOLO [18] is a bounding box predictor method for 2D cryoEM images, it is necessary to convert 3D tomograms into 2D slices and provide 2D annotations for each slice to train CrYOLO. For slicing the tomograms, we divided them into 2D x-y slices across the z-axis. If any macromolecule fell into the *z* = *t* th slice, we annotated all the 2D xy slices with z value in range [*t* − 12, *t* + 12] with that macromolecule in the same (x,y) coordinate as the radius of ribosome is maximally 12 voxels (12 × 1.348 = 16 nm) in the tomograms. For DeepETPicker [10], we used their publicly available codebase with the default settings for ribosome localization.

### Evaluation

For evaluation, we calculated the number of True Positives (TP), False Positives (FP), False Negatives (FN), Precision, Recall, and F1-score predicted by the baseline models, and our proposed models. To estimate the metrics, we used the annotations of ribosome coordinates provided in the original dataset as ground truth. If any predicted coordinate is within 10 voxels of Euclidean distance from a ground truth coordinate, it was regarded as a TP. Predicted coordinates not within 10 voxels were considered FP, while ground truth coordinates without nearby predicted coordinates were considered FN. Precision and Recall were calculated as: Precision 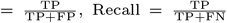. Finally, the F1 score was calculated as: 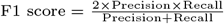.

By definition, precision penalizes FP but not FN. On the other hand, Recall penalizes FN, but not FP. Both FP and FN should be penalized for macromolecule localization. Consequently, we regard the F1 score as our primary evaluation criterion since it penalizes both.

### Experimental setup

In our experiments for TomoPicker, we used a batch size of 8 and an initial learning rate of 2 × 10^−3^, which has been reduced by a factor 0.5 if validation accuracy does not improve for 5 consecutive epochs. We trained TomoPicker and CrYOLO for 20 epochs, which we found to be sufficient for training convergence. However, we trained DeepETPicker for 100 epochs. We implemented our method in pytorch and trained the models using NVIDIA RTX A5000 GPUs.

We trained CrYOLO [18], DeepETPicker [10], and TomoPicker with three different losses (PN, PU, and KL) on the training tomogram. For VPP dataset, we used *N* ∈ {10, 100, 500} macromolecule coordinates from TS 0001 tomogram for training and *N* macromolecule coordinates from TS 0002 tomogram for validation while training the models. After training, the model was tested against all the tomograms. For training TomoPicker(PU) and TomoPicker(KL), we use the expected number of target macromolecules in the training tomogram, *E*_*p*_, as 2000, 2500, and 3000, where all gave similar results. In the final model, we used *E*_*p*_ = 2500. Though the training tomogram TS 0001 has 2450 ground truth macromolecules, we suppose the exact number of macromolecules is unknown in real-world scenarios, and only an estimate can be made. Consequently, we used this range of values to train the models and found that a rough estimate for *E*_*p*_ is sufficient to produce superior picking results. Moreover, such rough estimates can be readily provided by a structural biologist or even Large Vision-Language Models like GPT given the cellular volume the tomogram visualizes. We further observed that a much higher estimate maintains the performance (details in Supplementary document), whereas much lower value that expected count may hurt it. Consequently, we suggest the users to use a higher estimate than the expected count while training TomoPicker.

For the defocus-only dataset, we used *N* ∈ {10, 100, 500} macromolecule coordinates from the TS 029 tomogram for training and *N* macromolecule coordinates from the TS 030 tomogram for validation. After training, the model was tested against all the tomograms. For training TomoPicker(PU) and TomoPicker(KL), we use the expected number of macromolecules in the training tomogram, *E*_*p*_, as 2500, 3000, and 3500, where all gave almost identical results. In the final model, we used *E*_*p*_ = 3000.

For both the *S*.*Pombe* VPP and defocus-only datasets, the voxel spacing is 1.348 nm. Since the radius of the 80S ribosome macromolecule is 11 − 15 nm, we use 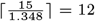 voxels as the macromolecule radius. For *M Pneumoniae* dataset, the voxel spacing is 1 nm. Since the radius of the 70S ribosome is 10 − 15 nm, we use 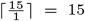 voxels as the macromolecule radius. This information is necessary for all the methods, including the baseline ones.

It takes approximately 0.006 seconds per epoch in a single GPU to train TomoPicker over a single subvolume of 32 × 32 × 32. For *S*.*Pombe* datasets, the training tomogram was split into 65,520 subvolumes of size 32 × 32 × 32. Hence, it took approximately 393 seconds, or 7 minutes, on a single GPU per epoch, resulting in 0.1 GPU-hours per epoch. For *M. Pneumoniae* dataset, the training tomogram was split into 4000 subvolumes of size 32 × 32 × 32. Consequently, each epoch required approximately 24 seconds on a single GPU, corresponding to 0.006 GPU-hours per epoch.

For inference, each tomogram in *S*.*Pombe* datasets took around 7 mins in a single GPU. On the other hand, each tomogram in *M. Pneumoniae* dataset took around only 30 seconds in a single GPU, due to their smaller sizes compared to Pombe tomograms.

### TomoPicker macromolecule localization closely match expert annotations using only 10 annotated macromolecule locations

We started our evaluation using the *S. Pombe* cellular tomogram dataset, which includes expert-annotated ground truth labels. We assessed whether TomoPicker could accurately predict macromolecule locations that closely match the ground truth annotations, even when trained using a few number of macromolecules. We performed experiments using annotated locations of *N* = 10 macromolecules in the training tomogram. This accounts for only 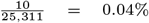 of the total macromolecules in the VPP dataset and 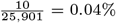 of the total macromolecules in the defocus-only dataset.

In the VPP dataset, we used the predicted and ground-truth macromolecule locations in the TS 0003 test tomogram and visualized them for a middle (*z* = 500) xy slice as bounding boxes. To convert the 3D predictions into 2D bounding boxes, we took slices across the z-axis and drew a bounding box around each predicted point (its coordinates *x*−*y*) in the corresponding xy slice. For each value on the z-axis (*z*), we also considered neighboring slices within a range [*z* − 12, *z* + 12], since the radius of the ribosome in these tomograms is maximally 12 voxels. Given the diameter of the ribosome of 24 voxels, we center the bounding boxes on the predicted points and size them to 24 voxels. We have provided a visualization for all model predictions and expert annotations as ground truth in Fig. 2.

From Fig. 2, it is evident that TomoPicker (KL) resembles the ground truth most closely. CrYOLO could not determine the locations for most macromolecular structures. In fact, in the middle xy slice of the TS 0003 tomogram, it detected only one macromolecular structure. However, DeepETPicker detected macromolecules almost randomly throughout the tomogram. This resulted in too many false positive detections. TomoPicker (PN), which simply regards all unlabeled voxels as non-macromolecule voxels during training, also detected a large number of macromolecules in regions without any actual macromolecule. TomoPicker (PU) offered much improvement over TomoPicker (PN) and provided results close to the ground truth. However, a large number of macromolecules have still been detected in regions without any actual macromolecules. TomoPicker (KL), on the other hand, provided the most plausible result, closely resembling the ground truth.

Similarly, in the defocus-only dataset, we used the predicted and ground-truth macromolecule locations in the TS 045 test tomogram and visualized them for a middle (*z* = 250) xy slice as bounding boxes. We have provided a visualization for all model predictions and expert annotations as ground truth in Fig. 3. Fig. 3 shows that CrYOLO, similar to its performance on VPP tomogram, could not determine the locations of most macromolecular structures. In fact, for TS 045, they provided several wrong detections. DeepETPicker, unlike its performance for the VPP tomogram, determined only a few macromolecules compared to the ground truth. TomoPicker (PN) determined several macromolecules, but also located many macromolecules in regions that did not contain any actual macromolecules. TomoPicker (PU) showed a slight improvement over TomoPicker (PN). Nevertheless, TomoPicker (KL), similar to the VPP dataset, most closely resembled the ground truth annotation.

### TomoPicker (KL) achieves almost comparable performance to supervised models with up to 98% less annotated data

Tables 1 and 2 report the F1 scores (rounded to two decimal places) for each method across individual tomograms and their average in the VPP and defocus-only datasets, respectively. Each table includes results for *N* = {10, 500} macromolecule training locations. Additional metrics and results for *N* = 100 are provided in the supplementary document.

**Table 1.**
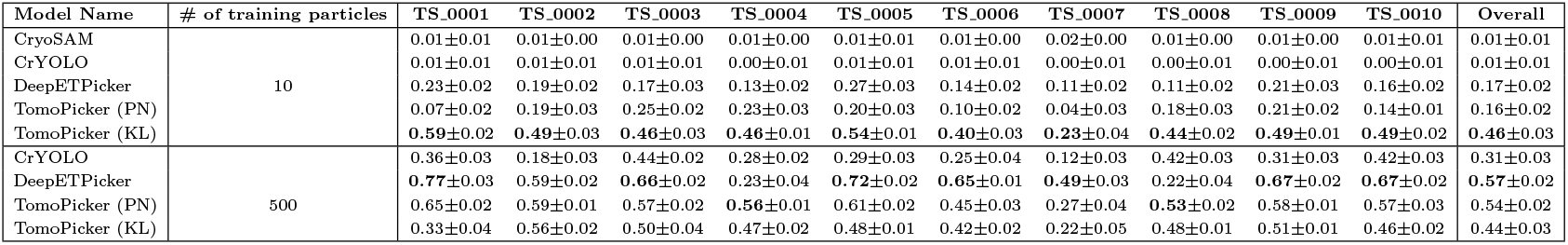
F1 score comparison across different methods on VPP *S. Pombe* cellular cryo-ET datasets. We trained for 3 independent random seeds and reported the mean ± std. dev. for each experimental setup. The precision and recall for these experiments, as well as the results for experiments with *N* = 100, are provided in the Supplementary Tables S1-S15.

| Model Name | # of training particles | TS_0001 | TS_0002 | TS_0003 | TS_0004 | TS_0005 | TS_0006 | TS_0007 | TS_0008 | TS_0009 | TS_0010 | Overall |
| --- | --- | --- | --- | --- | --- | --- | --- | --- | --- | --- | --- | --- |
| CryoSAM | 10 | 0.01±0.01 | 0.01±0.00 | 0.01±0.00 | 0.01±0.00 | 0.01±0.01 | 0.01±0.00 | 0.02±0.00 | 0.01±0.00 | 0.01±0.00 | 0.01±0.01 | 0.01±0.01 |
| CrYOLO | | 0.01±0.01 | 0.01±0.01 | 0.01±0.01 | 0.00±0.01 | 0.01±0.01 | 0.01±0.01 | 0.00±0.01 | 0.00±0.01 | 0.00±0.01 | 0.00±0.01 | 0.01±0.01 |
| DeepETPicker | | 0.23±0.02 | 0.19±0.02 | 0.17±0.03 | 0.13±0.02 | 0.27±0.03 | 0.14±0.02 | 0.11±0.02 | 0.11±0.02 | 0.21±0.03 | 0.16±0.02 | 0.17±0.02 |
| TomoPicker (PN) | | 0.07±0.02 | 0.19±0.03 | 0.25±0.02 | 0.23±0.03 | 0.20±0.03 | 0.10±0.02 | 0.04±0.03 | 0.18±0.03 | 0.21±0.02 | 0.14±0.01 | 0.16±0.02 |
| TomoPicker (KL) |  | **0.59±0.02** | **0.49±0.03** | **0.46±0.03** | **0.46±0.01** | **0.54±0.01** | **0.40±0.03** | **0.23±0.04** | **0.44±0.02** | **0.49±0.01** | **0.49±0.02** | **0.46±0.03** |
| CrYOLO | 500 | 0.36±0.03 | 0.18±0.03 | 0.44±0.02 | 0.28±0.02 | 0.29±0.03 | 0.25±0.04 | 0.12±0.03 | 0.42±0.03 | 0.31±0.03 | 0.42±0.03 | 0.31±0.03 |
| DeepETPicker | | **0.77±0.03** | 0.59±0.02 | **0.66±0.02** | 0.23±0.04 | **0.72±0.02** | **0.65±0.01** | **0.49±0.03** | 0.22±0.04 | **0.67±0.02** | **0.67±0.02** | **0.57±0.02** |
| TomoPicker (PN) | | 0.65±0.02 | 0.59±0.01 | 0.57±0.02 | **0.56±0.01** | 0.61±0.02 | 0.45±0.03 | 0.27±0.04 | **0.53±0.02** | 0.58±0.01 | 0.57±0.03 | 0.54±0.02 |
| TomoPicker (KL) | | 0.33±0.04 | 0.56±0.02 | 0.50±0.04 | 0.47±0.02 | 0.48±0.01 | 0.42±0.02 | 0.22±0.05 | 0.48±0.01 | 0.51±0.01 | 0.46±0.02 | 0.44±0.03 |

**Table 2.**
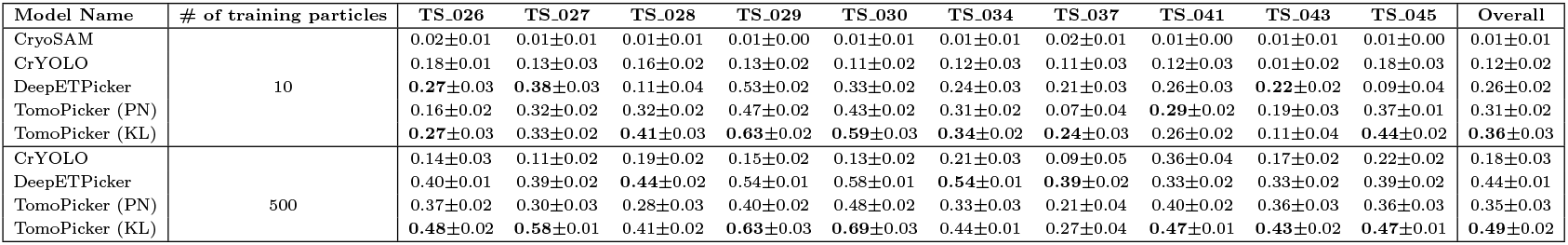
F1 score comparison across different methods on denoised defocus-only *S. Pombe* cryo-ET datasets. We trained for 3 independent random seeds and reported the mean ± std. dev. for each experimental setup. The precision and recall for these experiments, as well as the results for experiments with *N* = 100, are provided in the Supplementary Tables S16-S30.

We observe that the TomoPicker(KL) average F1 score for VPP and defocus datasets, while using only 10 annotated macromolecules for training, is almost the same as the supervised models trained with 500 annotated macromolecules. This suggests that TomoPicker (KL) achieves comparable performance with supervised models with 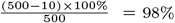 less annotated training data, suggesting a substantial reduction in the need for a burdensome manual annotation process for training learning-based macromolecule localization models.

Nevertheless, we observe that the performance of TomoPicker (KL) does not improve as markedly from *N* = 10 to *N* = 500 compared to fully supervised methods such as DeepETPicker, CrYOLO, and TomoPicker (PN). This is because TomoPicker (KL) is not solely driven by supervised signals; instead, it incorporates a strong bias from its KL-based positive–unlabeled learning loss. While this bias is highly beneficial under extremely sparse annotation conditions (e.g., *N* = 10), its advantage diminishes when sufficient supervised signal is available from larger annotation sets (e.g., *N* = 500). In such cases, methods trained purely on supervised data, such as TomoPicker (PN) or DeepETPicker, often achieve superior performance. Importantly, however, the design of TomoPicker (KL) is motivated by the scarcity of annotated data, a setting in which it demonstrates clear advantages.

### TomoPicker (KL) provides substantial improvement over supervised models under scarcity of annotated data

From Table 1 and Table 2, it is evident that TomoPicker provides a substantial improvement over existing models in the small number of annotated training data. When only 10 macromolecule locations are used to train it, TomoPicker (KL) provided an average F1 score of 0.46 in the VPP data set, which is ∼ 170% higher than the score obtained by the supervised model, DeepETPicker [10]. In Fig. 4, we plot the average F1 score achieved by each method for the VPP dataset with a varying number of training macromolecules. To better visualize the proportional relationship, we plot 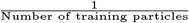 on the x-axis and the average F1 score by each method on the y-axis. The plot shows that with the increase of 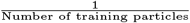 or decrease of Number of training particles, the F1 score improvement by TomoPicker (KL) over other models increases.

The defocus *S. Pombe* data set, unlike the VPP *S. Pombe* data set, contains more inter-tomogram variation. In addition, VPP images provide enhanced low frequency contrast due to the use of volta-phase plate, which improves signal-to-noise ratio (SNR) and facilitates voxel classification. On the other hand, defocus-only datasets have lower contrast and SNR. Since voxel classification inherently depends on contrast, it struggles to directly work on the defocus-only data. We performed denoising of the defocus-only tomograms before using voxel classifcation on the data to overcome this limitation. However, even after denoising, experiments on the defocus dataset show more irregular patterns in results compared to experiments on VPP, possibly due to issues related to denoising and high inter-tomogram variation between training and testing tomogram in the defocus-only dataset. Despite these facts, while training on only 10 annotations, TomoPicker (KL) achieved ∼ 40% improvement on the denoised defocus dataset over the supervised state-of-the-art DeepETPicker method.

We further assessed the performance of TomoPicker and the baseline methods under scarcity of annotated data on *Pneumoniae* cellular cryo-ET tomograms. Similar to our experiments on *S. Pombe* dataset, we used a single tomogram (with tomogram ID 00254) and used 10 annotated 70S ribosome locations to train the methods. Since we do not have the exact ground truth for these tomograms, quantitative evaluation would be less reliable. To this end, we qualitatively evaluated the macromolecule localization performance of TomoPicker and the supervised models. We provide some of the qualitative results in Figure 5. To make the qualitative results comprehensible, we performed difference of Gaussian (DoG) filtering on the tomograms before visualization. We provided visualizations of several slabs across different tomograms and corresponding inference results by TomoPicker (KL) and the most optimal supervised baseline DeepETPicker. The visualizations clearly demonstrate the superior performance of TomoPicker (KL) under sparse annotation training conditions. In fact, TomoPicker (KL) properly annotated macromolecule locations in cases where DeepETPicker performed totally random predictions. These results validate the generalizability of our claims across multiple species and cell types, encompassing diverse macromolecular compositions and varying tomogram voxel resolutions.

## Data Availability

The tomograms and the ground truth (expert annotations) macromolecule coordinates of *S. Pombe* cellular cryo-ET datasets (both VPP and defocus-only) all publicly available. They can be downloaded from EMPIAR-10988. They are also publicly available in the CZI (Chan-Zuckerberg Initiative) cryo-ET data portal as dataset ID 10000 (defocus-only) and 10001 (VPP). The tomograms of *M. Pneumoniae* cellular cryo-ET datasets are publicly available in EMPIAR-10499. The tomograms and the author provided coordinates are also available in the CZI (Chan-Zuckerberg Initiative) cryo-ET data portal as dataset ID 10003.

## Discussion & Conclusion

In this work, we have introduced a novel annotation-efficient macromolecule localization approach, TomoPicker, for 3D cellular cryo-ET images known as tomograms. In TomoPicker, we regarded macromolecule localization in 3D cryo-ET tomograms as a voxel classification problem. TomoPicker can be trained using very few macromolecule annotations in a single tomogram. To ensure effective localization in sparse annotation scenarios, we leveraged two different positive-unlabeled (PU) learning approaches to train the voxel classifier in TomoPicker. Whereas the previous methods for macromolecule localization were primarily evaluated on less crowded single-particle cryo-ET tomograms, we focused on highly crowded cellular cryo-ET tomograms. We trained and compared our method against the previous relevant methods on volta-phase-plate (VPP) and defocus-only *S. Pombe* cell cryo-ET tomograms and *M. Pneumoniae* cell cryo-ET tomograms. Our exhaustive experiments demonstrate the superior performance of TomoPicker over strong baseline methods when only 10 macromolecules, a minuscule portion of the total number of macromolecules in the tomogram dataset, are annotated for training. Ultimately, given these innovations, TomoPicker is poised to become a standard approach for macromolecule localization from cryo-ET images, within the complex intracellular environment of cells.

## Acknowledgement

This work was supported in part by U.S. NIH grants R01GM134020 and P41GM103712, NSF grants DBI-1949629, DBI-2238093, IIS-2007595, IIS-2211597, and MCB-2205148. This work was supported in part by Oracle Cloud credits and related resources provided by Oracle for Research, and the computational resources support from AMD HPC Fund. MRU were supported in part by a fellowship from CMU CMLH. This study was partially supported by UPMC Enterprises (ZF, MX).

