## Supplementary Information for "Localization of Macromolecules in Crowded Cellular Cryo-electron Tomograms from Extremely Sparse Labels"

### Supplementary Materials for Localization of Macromolecules in Crowded Cellular Cryo-electron Tomograms from Extremely Sparse Labels

This supplementary document contains further details on the related works and the extended tables of Table 1 and Table 2 in the main manuscript. The extended tables contain True Positive (TP), False Positive (FP), and False Negative (FN) particle counts, as well as precision, recall, and F1 scores obtained with TomoPicker, its variants, and the baseline methods on VPP and Defocus-only datasets.

#### 1 Related Works

**Template-Matching:** Before the development of learning-based models, Template Matching (TM) [3, 8] was the most used approach for macromolecule localization. In TM, structural templates from existing databases, such as the Protein Database (PDB) [2] and Electron Microscopy Data Bank (EMDB) [1], have usually been resolved through X-ray crystallography or single-particle cryo-EM. The structure of interest is first low-pass-filtered, and then randomly rotated in a fixed interval to create multiple templates. These templates are scanned throughout the cryo-ET tomograms in a sliding window manner, and a cross-correlation score is calculated between the templates and each subvolume. Thus, TM determines the location of the particles and their orientations with respect to the original template. Despite being widely used, this process is extremely time-consuming given the large size of the tomograms and the large number of templates to match with. Moreover, this is prone to template-dependent biases and cannot determine any structure without known existing templates. As a result, learning-based methods have been developed for macromolecule localization. The proposed method, TomoPicker, is also a learning-based picking method.

**Supervised learning-based methods:** To overcome the challenges of TM, several learning-based macromolecule localization methods have been developed. However, most of them [7, 4] are based on supervised learning, where a deep learning model is trained using a vast amount of manually annotated data. DeepFinder [7] and DeepiCT [4] use supervised UNet Networks to perform segmentation on cryo-ET tomograms, where predicted segmentation masks are used for macromolecule localization. Training these models requires manual annotation of segmentation masks. DeepETPicker [6] follows a similar strategy by placing Gaussian blobs in the picked location to create a ground truth segmentation map and trains a fully supervised UNet segmentation model. CrYOLO [9] performs supervised object detection using a YOLO-based object detection network. It requires the users to provide bounding box annotations instead of segmentation masks. However, CrYOLO [9] is originally developed for 2D single-particle cryo-EM images. Using this approach for 3D data requires performing 2D annotation on the 3D cryo-ET image slices and later converting the 2D predictions to 3D. Though providing boxes is easier than segmentation masks, the conversions between 2D and 3D are still a problem and successful training requires a large number of ground truth boxes to be manually annotated. Such manual annotation is also extremely time-consuming and burdensome. Unlike these methods, our method only requires providing the coordinates (approximately at the center) of a few particles to be annotated.

**Weakly supervised learning-based methods:** A few weakly-supervised learning-based macromolecule localization methods [5, 10] have been developed very recently that require only providing the center coordinates of a small number of particles in tomograms for training. These methods directly predict the center coordinate of particles given a whole 3D tomogram. However, as discussed earlier, they significantly downsample the tomograms to treat them as a single sample. Such downsampling increases the crowding of particles, which can be tolerable only for purified or a few prokaryote tomograms with sparsely located particles, but not for already crowded eukaryote tomograms. Moreover, they can be trained only on datasets containing many ($\geq 50$) similar tomograms. Unlike them, our method regards subvolumes from a tomogram as a sample and can be trained on a single tomogram. Since we do not require downsampling the tomograms, we do not introduce additional crowding in the tomograms, making our method suitable for crowded

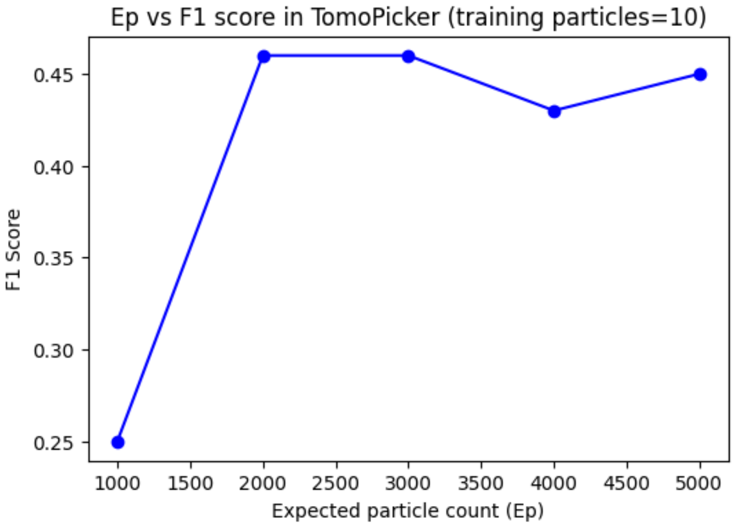


Figure S1: **Effect of "expected particle count in training tomogram" on the performance of TomoPicker (KL) in VPP S.Pombe cellular cryo-ET dataset under extremely sparse labels.** The actual count was 2450. We achieve consistent performance from Ep=2000 to Ep=5000.

eukaryote tomograms. In addition, our KL divergence-based PU learning method is methodologically much different from the PU learning method used in Huang et al. [5].

### 2 Training Details

#### 2.1 Choice of $E_p$

Both TomoPicker (PU) and TomoPicker (KL) requires a user provided expected macromolecule count for the training tomogram. We regard this value as $E_p$ which is used during training. We observed that an approximate value of $E_p$ is sufficient for our methods to perform well and an accurate estimate, which is hard to obtain, is not required. For instance, in the VPP S. Pombe dataset, we used TS_0001 as the training tomogram. The exact amount of 80S ribosomes in this tomogram was 2450. In our experiments of training TomoPicker (PU) and TomoPicker (KL), setting Ep=2000 or Ep=2500, or Ep=3000 in the VPP Pombe dataset resulted in similar performance. We further performed experiments with Ep=1000, Ep=4000, and Ep=5000. We found very similar performance with Ep=4000 and Ep=5000 to what we achieved with Ep=2500, indicating that setting a higher than expected particle count for Ep does not hurt the performance (Fig. S1). However, the performance deteriorated when Ep was set to 1000, suggesting that setting a value much lower than the expected particle count would hurt the performance.

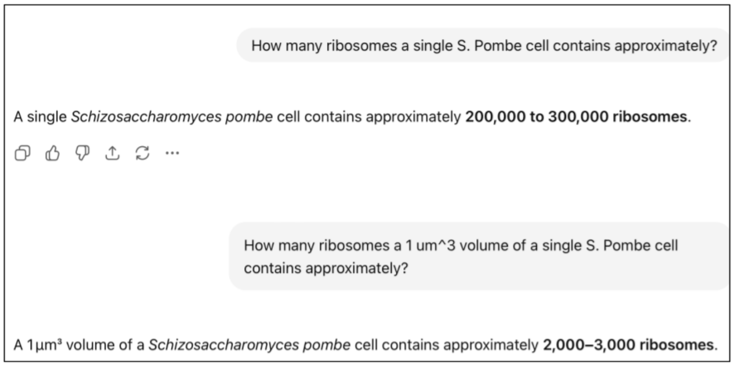


Figure S2: Estimating expected macromolecule count in training tomogram with GPT.

Furthermore, obtaining the approximate value for the expected particle count is often straightforward, given that we have information on the voxel spacing and thus cell size. For instance, the *S. Pombe* tomograms have a voxel spacing of 1.348 nm, and the dimensions of the VPP training tomogram was $500 \times 928 \times 960$.

This implies a tomogram would roughly capture 1 $\mu m^3$ of the S. Pombe cell. Since a full S. Pombe cell is about the size of 100 $\mu m^3$ and it contains approximately $2 \times 10^5$ to $3 \times 10^5$ ribosome particles, a 1 $\mu m^3$ volume of the cell contains around $2 \times 10^3$ to $3 \times 10^3$ 80S ribosome particles. Such information can be readily provided by biologists and even large-language models, such as GPT-5, GPT-4o (example in Fig. S2).

For *M. Pneumoniae* tomograms, the voxel spacing is 1 nm and the dimensions of the training tomogram is $306 \times 630 \times 630$. Consequently, the tomogram captures approximately 0.1 $\mu m^3$ of *M. Pneumoniae* cell. A single cell of *M. Pneumoniae* is also of approximate size 0.1 $\mu m^3$, suggesting the tomogram visualizes almost the whole cell. A single cell of *M. Pneumoniae* contains around 200 70S ribosomes, so does the training tomogram. However, given our consistent results with higher $E_p$, we set a high value of 400 while training TomoPicker on the *M. Pneumoniae* dataset.

#### 2.2 Loss Curves

We provide the training loss curves for TomoPicker (KL) on *S. Pombe* VPP and defocus datasets for different number of annotated training particles in Fig. S3.

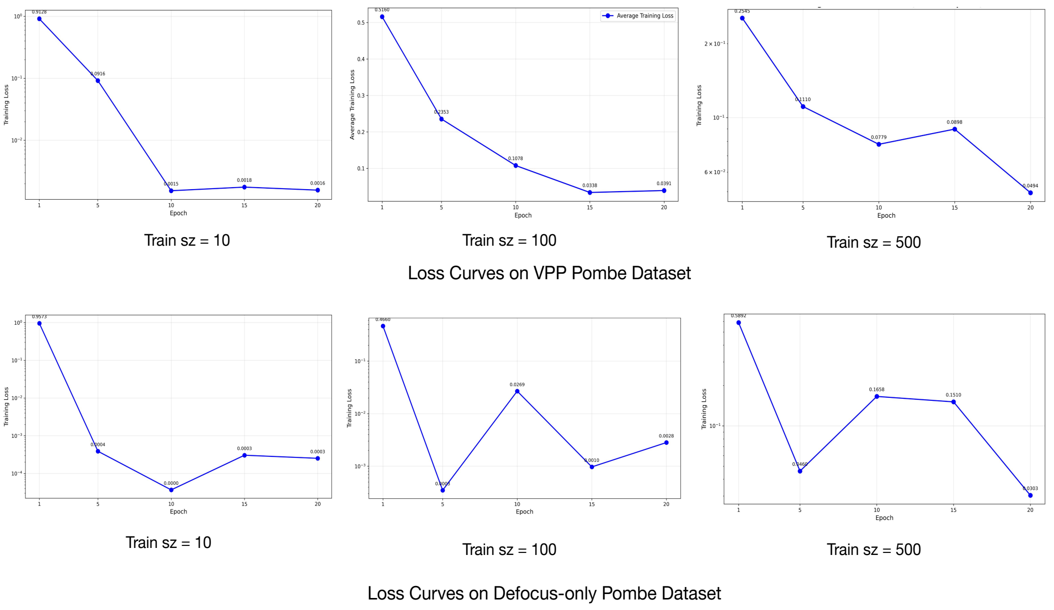


Figure S3: TomoPicker (KL) training loss curves

#### 2.3 Traditional Template Matching

The DeepETPicker [6] paper provided comparisons of DeepETPicker with template matching and DeepFinder across various datasets, where DeepETPicker consistently provided superior results. Consequently, we provided comparisons primarily with DeepETPicker, not DeepFinder [7] or traditional Template Matching. Moreover, in traditional template matching under sparse labels, one would require obtaining the template from the sparse annotation. In our extremely sparse scenario, we only used 10 annotations. Extracting subtomograms from only 10 locations and creating a template by averaging the 10 subtomograms with RELION [11] does not result in a template of sufficient resolution (Fig. S4). As a result, we believe traditional template matching under these extremely sparse label settings would not work well.

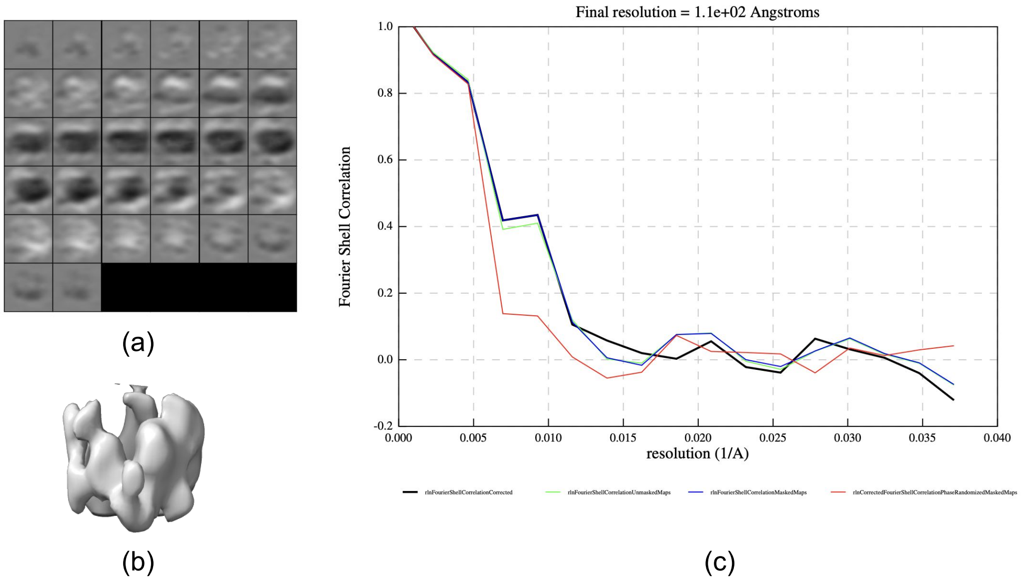


Figure S4: Template creation of 80S ribosome from 10 annotations on S.Pombe VPP tomograms using RELION [11]. (a) slice-by-slice visualization of the resulting template, (b) Iso-surface visualization of the resulting template, (c) FSC plot of subtomogram averaging with RELION.

### 3 VPP Ribosome Dataset

#### 3.1 Number of training particles=10

Table S1: Performance Metrics for CrYOLO

| Dataset | TP | FP | FN | Precision | Recall | F1 Score |
|---|---|---|---|---|---|---|
| TS_0001 | 10 | 5 | 2412 | 0.660000 | 0.005259 | 0.008000 |
| TS_0002 | 17 | 9 | 2325 | 0.653846 | 0.007259 | 0.014358 |
| TS_0003 | 7 | 10 | 2422 | 0.411765 | 0.002882 | 0.005724 |
| TS_0004 | 6 | 2 | 2961 | 0.750000 | 0.002022 | 0.004034 |
| TS_0005 | 13 | 8 | 3558 | 0.619048 | 0.003640 | 0.007238 |
| TS_0006 | 9 | 30 | 1327 | 0.230769 | 0.006737 | 0.013091 |
| TS_0007 | 1 | 8 | 616 | 0.043478 | 0.001621 | 0.003125 |
| TS_0008 | 2 | 0 | 2742 | 1.000000 | 0.000729 | 0.001457 |
| TS_0009 | 4 | 1 | 3478 | 0.800000 | 0.001149 | 0.002294 |
| TS_0010 | 6 | 0 | 3367 | 1.000000 | 0.001779 | 0.003551 |
| Overall | – | – | – | 0.616891 | 0.003282 | 0.006287 |

Table S2: Performance Metrics for DeepETPicker

| Dataset | TP | FP | FN | Precision | Recall | F1 Score |
|---|---|---|---|---|---|---|
| TS_0001 | 1424 | 8740 | 1026 | 0.14100232 | 0.58122449 | 0.22578087 |
| TS_0002 | 461 | 7181 | 1086 | 0.17793747 | 0.19684307 | 0.18730538 |
| TS_0003 | 535 | 3201 | 1894 | 0.14302128 | 0.22025542 | 0.17356042 |
| TS_0004 | 736 | 2147 | 2609 | 0.25510602 | 0.21904802 | 0.23587446 |
| TS_0005 | 1185 | 4109 | 2936 | 0.22383308 | 0.28735138 | 0.25174835 |
| TS_0006 | 415 | 4082 | 1981 | 0.09228375 | 0.17312381 | 0.12120347 |
| TS_0007 | 418 | 4081 | 2121 | 0.09211352 | 0.16420516 | 0.11866147 |
| TS_0008 | 402 | 4045 | 2342 | 0.09039801 | 0.14650148 | 0.11208642 |
| TS_0009 | 402 | 4072 | 2454 | 0.08911652 | 0.14048681 | 0.10989155 |
| TS_0010 | 999 | 4084 | 1801 | 0.19643863 | 0.35694339 | 0.25422937 |
| Overall = | – | – | – | 0.13521981 | 0.29208797 | 0.17093082 |

Table S3: Performance Metrics for TomoPicker (PN)

| Dataset | TP | FP | FN | Precision | Recall | F1 Score |
|---|---|---|---|---|---|---|
| TS_0001 | 145 | 1855 | 2305 | 0.0725 | 0.059184 | 0.065169 |
| TS_0002 | 421 | 1579 | 1921 | 0.2105 | 0.179761 | 0.19392 |
| TS_0003 | 560 | 1440 | 1869 | 0.2800 | 0.230548 | 0.252879 |
| TS_0004 | 680 | 2320 | 2287 | 0.226667 | 0.229188 | 0.227929 |
| TS_0005 | 657 | 2433 | 2914 | 0.2190 | 0.183892 | 0.1997 |
| TS_0006 | 145 | 1355 | 1191 | 0.096667 | 0.108533 | 0.102257 |
| TS_0007 | 23 | 477 | 594 | 0.0460 | 0.037277 | 0.041182 |
| TS_0008 | 503 | 2497 | 2241 | 0.167667 | 0.183139 | 0.175141 |
| TS_0009 | 688 | 2312 | 2296 | 0.229333 | 0.230678 | 0.230004 |
| TS_0010 | 436 | 2564 | 2937 | 0.145333 | 0.12962 | 0.13607 |
| Overall | – | – | – | 0.1693667 | 0.1538632 | 0.1607543 |

Table S4: Performance Metrics for TomoPicker (PU)

| Dataset | TP | FP | FN | Precision | Recall | F1 Score |
|---|---|---|---|---|---|---|
| TS_0001 | 681 | 1319 | 1769 | 0.3405 | 0.277959 | 0.306067 |
| TS_0002 | 347 | 1653 | 1795 | 0.1735 | 0.148164 | 0.159384 |
| TS_0003 | 265 | 1735 | 2125 | 0.1325 | 0.109098 | 0.119666 |
| TS_0004 | 473 | 2527 | 2494 | 0.157667 | 0.15942 | 0.158539 |
| TS_0005 | 2073 | 2437 | 2878 | 0.2310 | 0.194063 | 0.210927 |
| TS_0006 | 240 | 1260 | 1098 | 0.1600 | 0.179641 | 0.169252 |
| TS_0007 | 63 | 437 | 554 | 0.1260 | 0.102107 | 0.112802 |
| TS_0008 | 325 | 1620 | 1403 | 0.1625 | 0.188983 | 0.174604 |
| TS_0009 | 605 | 2483 | 2345 | 0.172333 | 0.148478 | 0.159519 |
| TS_0010 | 497 | 2503 | 2876 | 0.165667 | 0.147347 | 0.155971 |
| Overall | – | – | – | 0.17895 | 0.160877 | 0.1688719 |

Table S5: Performance Metrics for TomoPicker (GE-KL)

| Dataset | TP | FP | FN | Precision | Recall | F1 Score |
|---|---|---|---|---|---|---|
| TS_0001 | 1310 | 690 | 1140 | 0.655 | 0.534694 | 0.588764 |
| TS_0002 | 1060 | 940 | 1282 | 0.53 | 0.452605 | 0.488254 |
| TS_0003 | 1008 | 992 | 1421 | 0.504 | 0.414986 | 0.455182 |
| TS_0004 | 1360 | 1640 | 1607 | 0.453333 | 0.458375 | 0.45584 |
| TS_0005 | 1788 | 1212 | 1783 | 0.596 | 0.5007 | 0.544209 |
| TS_0006 | 568 | 932 | 768 | 0.378667 | 0.42515 | 0.400564 |
| TS_0007 | 370 | 487 | 206 | 0.432 | 0.210697 | 0.232766 |
| TS_0008 | 1253 | 1747 | 1491 | 0.417667 | 0.456633 | 0.436281 |
| TS_0009 | 1591 | 1409 | 1891 | 0.530333 | 0.456921 | 0.490898 |
| TS_0010 | 1569 | 1431 | 1804 | 0.523 | 0.465165 | 0.49239 |
| Overall | – | – | – | 0.4848 | 0.4375926 | 0.4585148 |

#### 3.2 Number of training particles=100

Table S6: Performance Metrics for CrYOLO

| Dataset | TP | FP | FN | Precision | Recall | F1 Score |
|---|---|---|---|---|---|---|
| TS_0001 | 1306 | 3322 | 1144 | 0.28 | 0.53 | 0.37 |
| TS_0002 | 555 | 1093 | 1787 | 0.34 | 0.24 | 0.28 |
| TS_0003 | 1377 | 5691 | 1052 | 0.19 | 0.57 | 0.29 |
| TS_0004 | 1677 | 7016 | 1290 | 0.19 | 0.57 | 0.29 |
| TS_0005 | 800 | 2167 | 2711 | 0.27 | 0.22 | 0.24 |
| TS_0006 | 210 | 1566 | 1126 | 0.12 | 0.16 | 0.13 |
| TS_0007 | 66 | 1133 | 551 | 0.06 | 0.11 | 0.07 |
| TS_0008 | 471 | 2273 | 2273 | 0.20 | 0.17 | 0.19 |
| TS_0009 | 1158 | 4372 | 2324 | 0.21 | 0.33 | 0.26 |
| TS_0010 | 1805 | 4289 | 1568 | 0.30 | 0.54 | 0.38 |
| Overall | – | – | – | 0.22 | 0.34 | 0.25 |

Table S7: Performance Metrics for DeepETPicker

| Dataset | TP | FP | FN | Precision | Recall | F1 Score |
|---|---|---|---|---|---|---|
| TS_0001 | 1632 | 506 | 818 | 0.76 | 0.67 | 0.70 |
| TS_0002 | 323 | 87 | 2019 | 0.79 | 0.14 | 0.23 |
| TS_0003 | 409 | 145 | 202 | 0.74 | 0.67 | 0.70 |
| TS_0004 | 79 | 32 | 2888 | 0.71 | 0.03 | 0.05 |
| TS_0005 | 1353 | 276 | 2218 | 0.83 | 0.38 | 0.52 |
| TS_0006 | 397 | 226 | 329 | 0.64 | 0.30 | 0.41 |
| TS_0007 | 302 | 417 | 315 | 0.42 | 0.49 | 0.45 |
| TS_0008 | 199 | 58 | 2545 | 0.77 | 0.07 | 0.13 |
| TS_0009 | 831 | 173 | 2651 | 0.83 | 0.24 | 0.37 |
| TS_0010 | 1023 | 299 | 2350 | 0.77 | 0.30 | 0.44 |
| Overall | – | – | – | 0.73 | 0.28 | 0.35 |

Table S8: Performance Metrics for TomoPicker (PN)

| Dataset | TP | FP | FN | Precision | Recall | F1 Score |
|---|---|---|---|---|---|---|
| TS_0001 | 1592 | 465 | 858 | 0.77 | 0.65 | 0.71 |
| TS_0002 | 51 | 46 | 1616 | 0.51 | 0.46 | 0.49 |
| TS_0003 | 594 | 1427 | 1835 | 0.41 | 0.38 | 0.39 |
| TS_0004 | 623 | 2397 | 2344 | 0.21 | 0.21 | 0.21 |
| TS_0005 | 1955 | 1108 | 1616 | 0.64 | 0.55 | 0.59 |
| TS_0006 | 235 | 780 | 1101 | 0.23 | 0.18 | 0.20 |
| TS_0007 | 247 | 771 | 370 | 0.24 | 0.40 | 0.30 |
| TS_0008 | 623 | 2385 | 2121 | 0.21 | 0.23 | 0.22 |
| TS_0009 | 1690 | 1344 | 1792 | 0.56 | 0.49 | 0.52 |
| TS_0010 | 1561 | 1490 | 1812 | 0.51 | 0.46 | 0.49 |
| Overall | – | – | – | 0.41 | 0.38 | 0.39 |

Table S9: Performance Metrics for TomoPicker (PU)

| Dataset | TP | FP | FN | Precision | Recall | F1 Score |
|---|---|---|---|---|---|---|
| TS_0001 | 1300 | 734 | 1150 | 0.64 | 0.53 | 0.58 |
| TS_0002 | 1049 | 1010 | 1293 | 0.51 | 0.45 | 0.48 |
| TS_0003 | 842 | 1188 | 1587 | 0.41 | 0.35 | 0.40 |
| TS_0004 | 1142 | 1892 | 1825 | 0.38 | 0.38 | 0.38 |
| TS_0005 | 1670 | 1378 | 1901 | 0.55 | 0.47 | 0.50 |
| TS_0006 | 293 | 718 | 1043 | 0.29 | 0.22 | 0.25 |
| TS_0007 | 218 | 789 | 399 | 0.35 | 0.33 | 0.27 |
| TS_0008 | 684 | 2326 | 2060 | 0.23 | 0.25 | 0.24 |
| TS_0009 | 1597 | 1440 | 1885 | 0.53 | 0.46 | 0.49 |
| TS_0010 | 1403 | 1645 | 1970 | 0.42 | 0.40 | 0.44 |
| Overall | – | – | – | 0.42 | 0.39 | 0.40 |

Table S10: Performance Metrics for TomoPicker (KL)

| Dataset | TP | FP | FN | Precision | Recall | F1 Score |
|---|---|---|---|---|---|---|
| TS_0001 | 1499 | 549 | 951 | 0.73 | 0.61 | 0.67 |
| TS_0002 | 1222 | 858 | 1120 | 0.59 | 0.52 | 0.55 |
| TS_0003 | 1162 | 890 | 1267 | 0.57 | 0.48 | 0.52 |
| TS_0004 | 484 | 1536 | 2483 | 0.24 | 0.16 | 0.19 |
| TS_0005 | 2074 | 1003 | 1497 | 0.67 | 0.58 | 0.62 |
| TS_0006 | 873 | 666 | 463 | 0.57 | 0.65 | 0.61 |
| TS_0007 | 254 | 758 | 363 | 0.25 | 0.41 | 0.32 |
| TS_0008 | 408 | 1601 | 2336 | 0.20 | 0.15 | 0.17 |
| TS_0009 | 675 | 1437 | 1867 | 0.33 | 0.19 | 0.25 |
| TS_0010 | 1879 | 1196 | 951 | 0.61 | 0.56 | 0.58 |
| Overall | – | – | – | 0.48 | 0.43 | 0.45 |

#### 3.3 Number of training particles=500

Table S11: Performance Metrics for CrYOLO

| Dataset | TP | FP | FN | Precision | Recall | F1 Score |
|---|---|---|---|---|---|---|
| TS_0001 | 1027 | 2154 | 1423 | 0.3229 | 0.4192 | 0.3648 |
| TS_0002 | 1604 | 13711 | 738 | 0.1047 | 0.6849 | 0.1817 |
| TS_0003 | 1347 | 2404 | 1082 | 0.3591 | 0.5545 | 0.4359 |
| TS_0004 | 1507 | 6439 | 1460 | 0.1897 | 0.5079 | 0.2762 |
| TS_0005 | 1841 | 7187 | 1730 | 0.2039 | 0.5155 | 0.2922 |
| TS_0006 | 733 | 3906 | 603 | 0.1580 | 0.5487 | 0.2454 |
| TS_0007 | 413 | 5811 | 204 | 0.0664 | 0.6694 | 0.1207 |
| TS_0008 | 1326 | 2203 | 1418 | 0.3654 | 0.4832 | 0.4161 |
| TS_0009 | 1608 | 5334 | 1874 | 0.2316 | 0.4618 | 0.3085 |
| TS_0010 | 1679 | 3036 | 1694 | 0.3561 | 0.4978 | 0.4152 |
| Overall | – | – | – | 0.23578 | 0.53429 | 0.30567 |

Table S12: Performance Metrics for DeepETPicker

| Dataset | TP | FP | FN | Precision | Recall | F1 Score |
|---|---|---|---|---|---|---|
| TS_0001 | 2206 | 1098 | 244 | 0.667676 | 0.900408 | 0.766771 |
| TS_0002 | 1052 | 193 | 1290 | 0.844498 | 0.449189 | 0.585653 |
| TS_0003 | 1367 | 334 | 1062 | 0.803645 | 0.562783 | 0.661985 |
| TS_0004 | 386 | 68 | 2581 | 0.85022 | 0.130098 | 0.225665 |
| TS_0005 | 2326 | 525 | 1245 | 0.815854 | 0.651358 | 0.723485 |
| TS_0006 | 840 | 395 | 496 | 0.680162 | 0.628743 | 0.653442 |
| TS_0007 | 450 | 783 | 167 | 0.364964 | 0.729335 | 0.486486 |
| TS_0008 | 353 | 81 | 2391 | 0.813364 | 0.128644 | 0.222152 |
| TS_0009 | 1951 | 411 | 1531 | 0.826 | 0.56031 | 0.667693 |
| TS_0010 | 1954 | 499 | 1419 | 0.796576 | 0.579306 | 0.670786 |
| Overall | – | – | – | 0.7463436 | 0.5320174 | 0.5665928 |

Table S13: Performance Metrics for TomoPicker (PN)

| Dataset | TP | FP | FN | Precision | Recall | F1 Score |
|---|---|---|---|---|---|---|
| TS_0001 | 1457 | 543 | 993 | 0.7285 | 0.594694 | 0.654831 |
| TS_0002 | 1285 | 715 | 1057 | 0.6425 | 0.548676 | 0.591893 |
| TS_0003 | 1260 | 740 | 1169 | 0.63 | 0.518732 | 0.568977 |
| TS_0004 | 1659 | 1341 | 1308 | 0.553 | 0.559151 | 0.556058 |
| TS_0005 | 2014 | 986 | 1557 | 0.671333 | 0.563988 | 0.612996 |
| TS_0006 | 637 | 863 | 699 | 0.424667 | 0.476796 | 0.449224 |
| TS_0007 | 149 | 351 | 468 | 0.298 | 0.241491 | 0.266786 |
| TS_0008 | 1511 | 1489 | 1233 | 0.503667 | 0.550656 | 0.526114 |
| TS_0009 | 1882 | 1118 | 1600 | 0.627333 | 0.540494 | 0.580685 |
| TS_0010 | 1810 | 1190 | 1563 | 0.603333 | 0.536614 | 0.568021 |
| Overall | – | – | – | 0.5682333 | 0.5131292 | 0.5375855 |

Table S14: Performance Metrics for TomoPicker (PU)

| Dataset | TP | FP | FN | Precision | Recall | F1 Score |
|---|---|---|---|---|---|---|
| TS_0001 | 1107 | 893 | 1343 | 0.5535 | 0.451837 | 0.497528 |
| TS_0002 | 1333 | 667 | 1009 | 0.6665 | 0.569172 | 0.614003 |
| TS_0003 | 1352 | 648 | 1077 | 0.676 | 0.556608 | 0.610522 |
| TS_0004 | 1597 | 1403 | 1370 | 0.532333 | 0.538254 | 0.535277 |
| TS_0005 | 1855 | 1145 | 1716 | 0.618333 | 0.519462 | 0.564602 |
| TS_0006 | 502 | 998 | 834 | 0.334667 | 0.375749 | 0.35402 |
| TS_0007 | 53 | 447 | 564 | 0.106 | 0.0859 | 0.094897 |
| TS_0008 | 1367 | 1633 | 1377 | 0.455667 | 0.498178 | 0.475975 |
| TS_0009 | 1862 | 1138 | 1620 | 0.620667 | 0.53475 | 0.574514 |
| TS_0010 | 1709 | 1291 | 1664 | 0.569667 | 0.506671 | 0.536325 |
| Overall | – | – | – | 0.5133334 | 0.4636581 | 0.4857663 |

Table S15: Performance Metrics for TomoPicker (GE-KL)

| Dataset | TP | FP | FN | Precision | Recall | F1 Score |
|---|---|---|---|---|---|---|
| TS_0001 | 733 | 1267 | 1717 | 0.3665 | 0.299184 | 0.329438 |
| TS_0002 | 1223 | 777 | 1119 | 0.6115 | 0.522203 | 0.563335 |
| TS_0003 | 1104 | 896 | 1325 | 0.552 | 0.454508 | 0.498532 |
| TS_0004 | 1417 | 1583 | 1500 | 0.472333 | 0.477587 | 0.474946 |
| TS_0005 | 1588 | 1412 | 1983 | 0.529333 | 0.444693 | 0.483366 |
| TS_0006 | 596 | 904 | 740 | 0.397333 | 0.446108 | 0.42031 |
| TS_0007 | 125 | 375 | 492 | 0.25 | 0.202593 | 0.223814 |
| TS_0008 | 1390 | 1610 | 1354 | 0.463333 | 0.50656 | 0.483983 |
| TS_0009 | 1667 | 1333 | 1815 | 0.555667 | 0.478748 | 0.513447 |
| TS_0010 | 1454 | 1546 | 1919 | 0.484667 | 0.43107 | 0.4563 |
| Overall | – | – | – | 0.4682666 | 0.4263254 | 0.4448341 |

### 4 Defocus Ribosome Performances

#### 4.1 Number of training particles=10

Table S16: Performance Metrics for CrYOLO

| Dataset | TP | FP | FN | Precision | Recall | F1 Score |
|---|---|---|---|---|---|---|
| TS_026 | 290 | 212 | 2365 | 0.570 | 0.1092 | 0.183 |
| TS_027 | 139 | 278 | 1534 | 0.33333 | 0.08308 | 0.13301 |
| TS_028 | 506 | 468 | 4799 | 0.51951 | 0.09538 | 0.16117 |
| TS_029 | 227 | 499 | 2670 | 0.31267 | 0.07836 | 0.12531 |
| TS_030 | 182 | 249 | 2601 | 0.42227 | 0.0654 | 0.11325 |
| TS_034 | 252 | 202 | 3531 | 0.55507 | 0.06661 | 0.11895 |
| TS_037 | 112 | 260 | 1534 | 0.30108 | 0.06804 | 0.111 |
| TS_041 | 176 | 34 | 2637 | 0.8381 | 0.06257 | 0.11644 |
| TS_043 | 7 | 46 | 1808 | 0.13208 | 0.00386 | 0.00749 |
| TS_045 | 245 | 193 | 2103 | 0.55936 | 0.10434 | 0.17588 |
| Overall | – | – | – | 0.45435 | 0.07368 | 0.12455 |

Table S17: Performance Metrics for DeepETPicker

| Dataset | TP | FP | FN | Precision | Recall | F1 Score |
|---|---|---|---|---|---|---|
| TS_026 | 231 | 618 | 607 | 0.272085 | 0.275656 | 0.273859 |
| TS_027 | 884 | 2103 | 789 | 0.295949 | 0.528392 | 0.379399 |
| TS_028 | 310 | 83 | 4995 | 0.788804 | 0.058435 | 0.10881 |
| TS_029 | 1597 | 1571 | 1300 | 0.504104 | 0.551126 | 0.526628 |
| TS_030 | 625 | 329 | 2158 | 0.655136 | 0.224578 | 0.334493 |
| TS_034 | 566 | 307 | 3217 | 0.648339 | 0.149617 | 0.243127 |
| TS_037 | 216 | 168 | 1430 | 0.5625 | 0.131227 | 0.212808 |
| TS_041 | 493 | 527 | 2320 | 0.483333 | 0.175528 | 0.25724 |
| TS_043 | 376 | 1176 | 1439 | 0.242268 | 0.207163 | 0.223344 |
| TS_045 | 118 | 89 | 2230 | 0.570048 | 0.050256 | 0.092368 |
| Overall | – | – | – | 0.5022566 | 0.2351842 | 0.2652076 |

Table S18: Performance Metrics for TomoPicker (PN)

| Dataset | TP | FP | FN | Precision | Recall | F1 Score |
|---|---|---|---|---|---|---|
| TS_026 | 223 | 1777 | 615 | 0.1115 | 0.26611 | 0.157153 |
| TS_027 | 592 | 1408 | 1081 | 0.296 | 0.353855 | 0.322352 |
| TS_028 | 1184 | 816 | 4121 | 0.592 | 0.223186 | 0.324162 |
| TS_029 | 1375 | 1625 | 1522 | 0.458333 | 0.474629 | 0.466339 |
| TS_030 | 1243 | 1757 | 1540 | 0.414333 | 0.44664 | 0.429881 |
| TS_034 | 831 | 669 | 2952 | 0.554 | 0.219667 | 0.314594 |
| TS_037 | 77 | 423 | 1569 | 0.154 | 0.04678 | 0.071761 |
| TS_041 | 841 | 2159 | 1972 | 0.280333 | 0.299869 | 0.289351 |
| TS_043 | 446 | 2554 | 1369 | 0.148667 | 0.245733 | 0.185254 |
| TS_045 | 979 | 2021 | 1369 | 0.326333 | 0.416951 | 0.366118 |
| Overall | – | – | – | 0.3335499 | 0.2992517 | 0.3077569 |

Table S19: Performance Metrics for TomoPicker (PU)

| Dataset | TP | FP | FN | Precision | Recall | F1 Score |
|---|---|---|---|---|---|---|
| TS_026 | 245 | 1755 | 593 | 0.1225 | 0.292363 | 0.172657 |
| TS_027 | 365 | 1635 | 1308 | 0.1825 | 0.218171 | 0.198748 |
| TS_028 | 1099 | 901 | 4206 | 0.5495 | 0.207163 | 0.30089 |
| TS_029 | 836 | 2164 | 2061 | 0.278667 | 0.288574 | 0.283534 |
| TS_030 | 1242 | 1758 | 1541 | 0.414 | 0.446821 | 0.429535 |
| TS_034 | 598 | 902 | 3185 | 0.398667 | 0.158076 | 0.226387 |
| TS_037 | 133 | 367 | 1513 | 0.266 | 0.080802 | 0.123952 |
| TS_041 | 624 | 2376 | 2189 | 0.208 | 0.221827 | 0.214691 |
| TS_043 | 274 | 2726 | 1541 | 0.091333 | 0.150964 | 0.113811 |
| TS_045 | 826 | 2174 | 1522 | 0.275333 | 0.351789 | 0.308901 |
| Overall | – | – | – | 0.278035 | 0.241755 | 0.2373106 |

Table S20: Performance Metrics for TomoPicker (GE-KL)

| Dataset | TP | FP | FN | Precision | Recall | F1 Score |
|---|---|---|---|---|---|---|
| TS_026 | 380 | 1620 | 458 | 0.190 | 0.453461 | 0.267794 |
| TS_027 | 606 | 1394 | 1067 | 0.303 | 0.362224 | 0.329957 |
| TS_028 | 1510 | 490 | 3795 | 0.755 | 0.284637 | 0.413415 |
| TS_029 | 1849 | 1151 | 1048 | 0.616333 | 0.638246 | 0.627099 |
| TS_030 | 1709 | 1291 | 1074 | 0.569667 | 0.614086 | 0.591043 |
| TS_034 | 886 | 614 | 2897 | 0.590667 | 0.234206 | 0.335415 |
| TS_037 | 256 | 224 | 1390 | 0.512 | 0.155529 | 0.238583 |
| TS_041 | 758 | 2242 | 2055 | 0.252667 | 0.269463 | 0.260795 |
| TS_043 | 275 | 2725 | 1540 | 0.091667 | 0.151515 | 0.114226 |
| TS_045 | 1169 | 1831 | 1179 | 0.389667 | 0.497871 | 0.437173 |
| Overall | – | – | – | 0.4270668 | 0.3661238 | 0.3615518 |

#### 4.2 Number of training particles=100

Table S21: Performance Metrics for CrYOLO

| Dataset | TP | FP | FN | Precision | Recall | F1 Score |
|---|---|---|---|---|---|---|
| TS_026 | 1520 | 4119 | 679 | 0.2600 | 0.6900 | 0.3800 |
| TS_027 | 790 | 3908 | 883 | 0.1682 | 0.4722 | 0.2480 |
| TS_028 | 3090 | 7250 | 2215 | 0.2988 | 0.5825 | 0.3950 |
| TS_029 | 1485 | 6157 | 1412 | 0.1943 | 0.5126 | 0.2818 |
| TS_030 | 1566 | 5686 | 1217 | 0.2159 | 0.5627 | 0.3121 |
| TS_034 | 1935 | 3798 | 1848 | 0.3375 | 0.5115 | 0.4067 |
| TS_037 | 892 | 6259 | 754 | 0.1247 | 0.5419 | 0.2028 |
| TS_041 | 1201 | 1588 | 1612 | 0.4306 | 0.4269 | 0.4288 |
| TS_043 | 179 | 888 | 1636 | 0.1678 | 0.0986 | 0.1242 |
| TS_045 | 1323 | 3644 | 1025 | 0.2664 | 0.5635 | 0.3617 |
| Overall | – | – | – | 0.24642 | 0.49624 | 0.31411 |

Table S22: Performance Metrics for DeepETPciker

| Dataset | TP | FP | FN | Precision | Recall | F1 Score |
|---|---|---|---|---|---|---|
| TS_026 | 227 | 357 | 611 | 0.3887 | 0.2709 | 0.3193 |
| TS_027 | 442 | 523 | 1231 | 0.4580 | 0.2642 | 0.3351 |
| TS_028 | 601 | 112 | 4704 | 0.8429 | 0.1133 | 0.1997 |
| TS_029 | 853 | 395 | 2044 | 0.6835 | 0.2944 | 0.4116 |
| TS_030 | 593 | 187 | 2190 | 0.7603 | 0.2131 | 0.3329 |
| TS_034 | 707 | 182 | 3076 | 0.7953 | 0.1869 | 0.3200 |
| TS_037 | 270 | 198 | 1376 | 0.5769 | 0.1640 | 0.2554 |
| TS_041 | 288 | 217 | 2525 | 0.5703 | 0.1024 | 0.1736 |
| TS_043 | 241 | 386 | 1574 | 0.3844 | 0.1327 | 0.1980 |
| TS_045 | 251 | 87 | 2097 | 0.7426 | 0.1069 | 0.1869 |
| Overall | – | – | – | 0.6203 | 0.1849 | 0.2715 |

Table S23: Performance Metrics for TomoPicker (PN)

| Dataset | TP | FP | FN | Precision | Recall | F1 Score |
|---|---|---|---|---|---|---|
| TS_026 | 404 | 1596 | 434 | 0.202 | 0.4821 | 0.284708 |
| TS_027 | 728 | 1272 | 945 | 0.364 | 0.435146 | 0.396406 |
| TS_028 | 1156 | 844 | 4149 | 0.578 | 0.217908 | 0.316496 |
| TS_029 | 1906 | 1094 | 991 | 0.635333 | 0.657922 | 0.646434 |
| TS_030 | 1684 | 1316 | 1099 | 0.561333 | 0.605102 | 0.582397 |
| TS_034 | 897 | 603 | 2886 | 0.598 | 0.237113 | 0.33958 |
| TS_037 | 211 | 289 | 1435 | 0.422 | 0.12819 | 0.196645 |
| TS_041 | 599 | 2401 | 2214 | 0.199667 | 0.21294 | 0.20609 |
| TS_043 | 256 | 2744 | 1559 | 0.085333 | 0.141047 | 0.106334 |
| TS_045 | 878 | 2122 | 1470 | 0.292667 | 0.373935 | 0.328347 |
| Overall | – | – | – | 0.3938333 | 0.3491403 | 0.3403433 |

Table S24: Performance Metrics for TomoPicker (PU)

| Dataset | TP | FP | FN | Precision | Recall | F1 Score |
|---|---|---|---|---|---|---|
| TS_026 | 280 | 1720 | 558 | 0.14 | 0.334129 | 0.197322 |
| TS_027 | 542 | 1458 | 1131 | 0.271 | 0.323969 | 0.295127 |
| TS_028 | 1341 | 659 | 3964 | 0.6705 | 0.25278 | 0.367146 |
| TS_029 | 1775 | 1225 | 1122 | 0.591667 | 0.612703 | 0.602001 |
| TS_030 | 1658 | 1342 | 1125 | 0.552667 | 0.59576 | 0.573405 |
| TS_034 | 768 | 732 | 3015 | 0.512 | 0.203013 | 0.290744 |
| TS_037 | 196 | 304 | 1450 | 0.392 | 0.119077 | 0.182665 |
| TS_041 | 504 | 2496 | 2309 | 0.168 | 0.179168 | 0.173404 |
| TS_043 | 180 | 2820 | 1635 | 0.06 | 0.099174 | 0.074766 |
| TS_045 | 1109 | 1891 | 1239 | 0.369667 | 0.472317 | 0.414734 |
| Overall | – | – | – | 0.3727501 | 0.319209 | 0.3171314 |

Table S25: Performance Metrics for TomoPicker (KL)

| Dataset | TP | FP | FN | Precision | Recall | F1 Score |
|---|---|---|---|---|---|---|
| TS_026 | 203 | 1797 | 635 | 0.1015 | 0.242243 | 0.143058 |
| TS_027 | 451 | 1549 | 1222 | 0.2255 | 0.269576 | 0.245576 |
| TS_028 | 1385 | 615 | 3920 | 0.6925 | 0.261074 | 0.379192 |
| TS_029 | 1761 | 1239 | 1136 | 0.587 | 0.60787 | 0.597253 |
| TS_030 | 1588 | 1412 | 1195 | 0.529333 | 0.570607 | 0.549196 |
| TS_034 | 706 | 794 | 3077 | 0.470667 | 0.186624 | 0.267727 |
| TS_037 | 180 | 320 | 1466 | 0.36 | 0.109356 | 0.167754 |
| TS_041 | 417 | 2583 | 2396 | 0.139 | 0.14824 | 0.143472 |
| TS_043 | 165 | 2835 | 1650 | 0.055 | 0.090909 | 0.068536 |
| TS_045 | 1063 | 1937 | 1285 | 0.354333 | 0.452726 | 0.397532 |
| Overall | – | – | – | 0.2958841 | 0.2892325 | 0.2875034 |

#### 4.3 Number of training particles=500

Table S26: Performance Metrics for CrYOLO

| Dataset | TP | FP | FN | Precision | Recall | F1 Score |
|---|---|---|---|---|---|---|
| TS_026 | 1322 | 26610 | 156 | 0.0800 | 0.8900 | 0.1400 |
| TS_027 | 1506 | 24315 | 167 | 0.0583 | 0.9002 | 0.1096 |
| TS_028 | 4933 | 41740 | 372 | 0.1057 | 0.9299 | 0.1898 |
| TS_029 | 2632 | 30634 | 265 | 0.0791 | 0.9085 | 0.1456 |
| TS_030 | 2574 | 33215 | 209 | 0.0719 | 0.9249 | 0.1335 |
| TS_034 | 3468 | 25292 | 315 | 0.1206 | 0.9167 | 0.2131 |
| TS_037 | 1456 | 28264 | 190 | 0.0490 | 0.8846 | 0.0928 |
| TS_041 | 1396 | 7924 | 494 | 0.2264 | 0.8244 | 0.3552 |
| TS_043 | 977 | 8615 | 838 | 0.1018 | 0.5383 | 0.1713 |
| TS_045 | 2117 | 14810 | 231 | 0.1251 | 0.9016 | 0.2197 |
| Overall | – | – | – | 0.1018 | 0.86191 | 0.17706 |

Table S27: Performance Metrics for DeepETPicker

| Dataset | TP | FP | FN | Precision | Recall | F1 Score |
|---|---|---|---|---|---|---|
| TS_026 | 512 | 1238 | 326 | 0.292571 | 0.610979 | 0.395672 |
| TS_027 | 891 | 1952 | 782 | 0.313401 | 0.532756 | 0.394597 |
| TS_028 | 1630 | 522 | 3675 | 0.757435 | 0.307257 | 0.437173 |
| TS_029 | 1618 | 1502 | 1279 | 0.518595 | 0.558509 | 0.53781 |
| TS_030 | 1436 | 733 | 1347 | 0.662056 | 0.515999 | 0.579968 |
| TS_034 | 1628 | 648 | 2155 | 0.71529 | 0.430346 | 0.537382 |
| TS_037 | 637 | 979 | 1009 | 0.394138 | 0.386999 | 0.390558 |
| TS_041 | 670 | 518 | 2143 | 0.563973 | 0.238118 | 0.334916 |
| TS_043 | 600 | 1247 | 1215 | 0.324851 | 0.330579 | 0.327669 |
| TS_045 | 726 | 630 | 1622 | 0.535398 | 0.309199 | 0.392009 |
| Overall | – | – | – | 0.5077748 | 0.4220614 | 0.432778 |

Table S28: Performance Metrics for TomoPicker (PN)

| Dataset | TP | FP | FN | Precision | Recall | F1 Score |
|---|---|---|---|---|---|---|
| TS_026 | 530 | 1470 | 308 | 0.265 | 0.632458 | 0.373502 |
| TS_027 | 555 | 1445 | 1118 | 0.2775 | 0.331739 | 0.302025 |
| TS_028 | 1029 | 971 | 4276 | 0.5145 | 0.193968 | 0.281725 |
| TS_029 | 1183 | 1817 | 1714 | 0.394333 | 0.408353 | 0.401221 |
| TS_030 | 1388 | 1612 | 1395 | 0.462667 | 0.498742 | 0.480082 |
| TS_034 | 876 | 624 | 2907 | 0.584 | 0.231562 | 0.33163 |
| TS_037 | 221 | 279 | 1425 | 0.442 | 0.134265 | 0.205965 |
| TS_041 | 1166 | 1834 | 1647 | 0.388667 | 0.414507 | 0.40117 |
| TS_043 | 873 | 2127 | 942 | 0.291 | 0.480992 | 0.362617 |
| TS_045 | 963 | 2037 | 1385 | 0.321 | 0.410136 | 0.360135 |
| Overall | – | – | – | 0.3940667 | 0.3736719 | 0.3500198 |

Table S29: Performance Metrics for TomoPicker (PU)

| Dataset | TP | FP | FN | Precision | Recall | F1 Score |
|---|---|---|---|---|---|---|
| TS_026 | 517 | 1483 | 321 | 0.2585 | 0.616945 | 0.364341 |
| TS_027 | 614 | 1386 | 1059 | 0.307 | 0.367005 | 0.334432 |
| TS_028 | 1109 | 891 | 4196 | 0.5545 | 0.209048 | 0.303628 |
| TS_029 | 1076 | 1924 | 1821 | 0.358667 | 0.371149 | 0.364931 |
| TS_030 | 1467 | 1533 | 1316 | 0.489 | 0.527129 | 0.507349 |
| TS_034 | 937 | 563 | 2846 | 0.624667 | 0.247687 | 0.354723 |
| TS_037 | 249 | 251 | 1397 | 0.498 | 0.151276 | 0.232065 |
| TS_041 | 1183 | 1817 | 1630 | 0.394333 | 0.420547 | 0.407019 |
| TS_043 | 952 | 2048 | 863 | 0.317333 | 0.524518 | 0.395431 |
| TS_045 | 980 | 2020 | 1368 | 0.326667 | 0.417376 | 0.366492 |
| Overall | – | – | – | 0.4128667 | 0.385295 | 0.3630306 |

Table S30: Performance Metrics for TomoPicker (KL)

| Dataset | TP | FP | FN | Precision | Recall | F1 Score |
|---|---|---|---|---|---|---|
| TS_026 | 685 | 1315 | 153 | 0.3425 | 0.817422 | 0.482734 |
| TS_027 | 1059 | 941 | 614 | 0.5295 | 0.632995 | 0.57664 |
| TS_028 | 1506 | 494 | 3799 | 0.753 | 0.283883 | 0.41232 |
| TS_029 | 1867 | 1133 | 1030 | 0.622333 | 0.64446 | 0.633023 |
| TS_030 | 1992 | 1008 | 791 | 0.664 | 0.715774 | 0.689816 |
| TS_034 | 1158 | 342 | 2625 | 0.772 | 0.306106 | 0.438387 |
| TS_037 | 289 | 211 | 1357 | 0.578 | 0.175577 | 0.269338 |
| TS_041 | 1363 | 1637 | 1450 | 0.453333 | 0.484356 | 0.468949 |
| TS_043 | 1046 | 1954 | 769 | 0.348667 | 0.576309 | 0.434476 |
| TS_045 | 1265 | 1735 | 1083 | 0.421667 | 0.538756 | 0.473074 |
| Overall | – | – | – | 0.5486 | 0.5175818 | 0.4878037 |

### References


[1] Emdb—the electron microscopy data bank. *Nucleic acids research*, 52(D1):D456–D465, 2024.

[2] Helen Berman, Kim Henrick, and Haruki Nakamura. Announcing the worldwide protein data bank. *Nature structural & molecular biology*, 10(12):980–980, 2003.

[3] Jochen Böhm, Achilleas S Frangakis, Reiner Hegerl, Stephan Nickell, Dieter Typke, and Wolfgang Baumeister. Toward detecting and identifying macromolecules in a cellular context: template matching applied to electron tomograms. *Proceedings of the National Academy of Sciences*, 97(26):14245–14250, 2000.

[4] Irene de Teresa-Trueba, Sara K Goetz, Alexander Mattausch, Frosina Stojanovska, Christian E Zimmerli, Mauricio Toro-Nahuelpan, Dorothy WC Cheng, Fergus Tollervey, Constantin Pape, Martin Beck,

et al. Convolutional networks for supervised mining of molecular patterns within cellular context. *Nature Methods*, 20(2):284–294, 2023.
